# MTG16 (CBFA2T3) regulates colonic epithelial differentiation, colitis, and tumorigenesis by repressing E protein transcription factors

**DOI:** 10.1101/2021.11.03.467178

**Authors:** Rachel E. Brown, Justin Jacobse, Shruti A. Anant, Koral M. Blunt, Bob Chen, Paige N. Vega, Chase T. Jones, Jennifer M. Pilat, Frank Revetta, Aidan H. Gorby, Kristy R. Stengel, Yash A. Choksi, Kimmo Palin, M. Blanca Piazuelo, M. Kay Washington, Ken S. Lau, Jeremy A. Goettel, Scott W. Hiebert, Sarah P. Short, Christopher S. Williams

## Abstract

Aberrant epithelial differentiation and regeneration contribute to colon pathologies including inflammatory bowel disease (IBD) and colitis-associated cancer (CAC). MTG16 (CBFA2T3) is a transcriptional corepressor expressed in the colonic epithelium. MTG16 deficiency in mice exacerbates colitis and increases tumor burden in CAC, though the underlying mechanisms remain unclear. Here, we identified MTG16 as a central mediator of epithelial differentiation, promoting goblet and restraining enteroendocrine cell development in homeostasis and enabling regeneration following dextran sulfate sodium (DSS)-induced colitis. Transcriptomic analyses implicated increased E box-binding transcription factor (E protein) activity in MTG16-deficient colon crypts. Using a novel mouse model with a point mutation that disrupts MTG16:E protein complex formation (*Mtg16^P209T^*), we established that MTG16 exerts control over colonic epithelial differentiation and regeneration by repressing E protein-mediated transcription. Mimicking murine colitis, *MTG16* expression was increased in biopsies from patients with active IBD compared to unaffected controls. Finally, uncoupling MTG16:E protein interactions only partially phenocopied the enhanced tumorigenicity of *Mtg16^-/-^* colon in the azoxymethane(AOM)/DSS-induced model of CAC, indicating that MTG16 protects from tumorigenesis through additional mechanisms. Collectively, our results demonstrate that MTG16, via its repression of E protein targets, is a key regulator of cell fate decisions during colon homeostasis, colitis, and cancer.

**Graphical Abstract:** 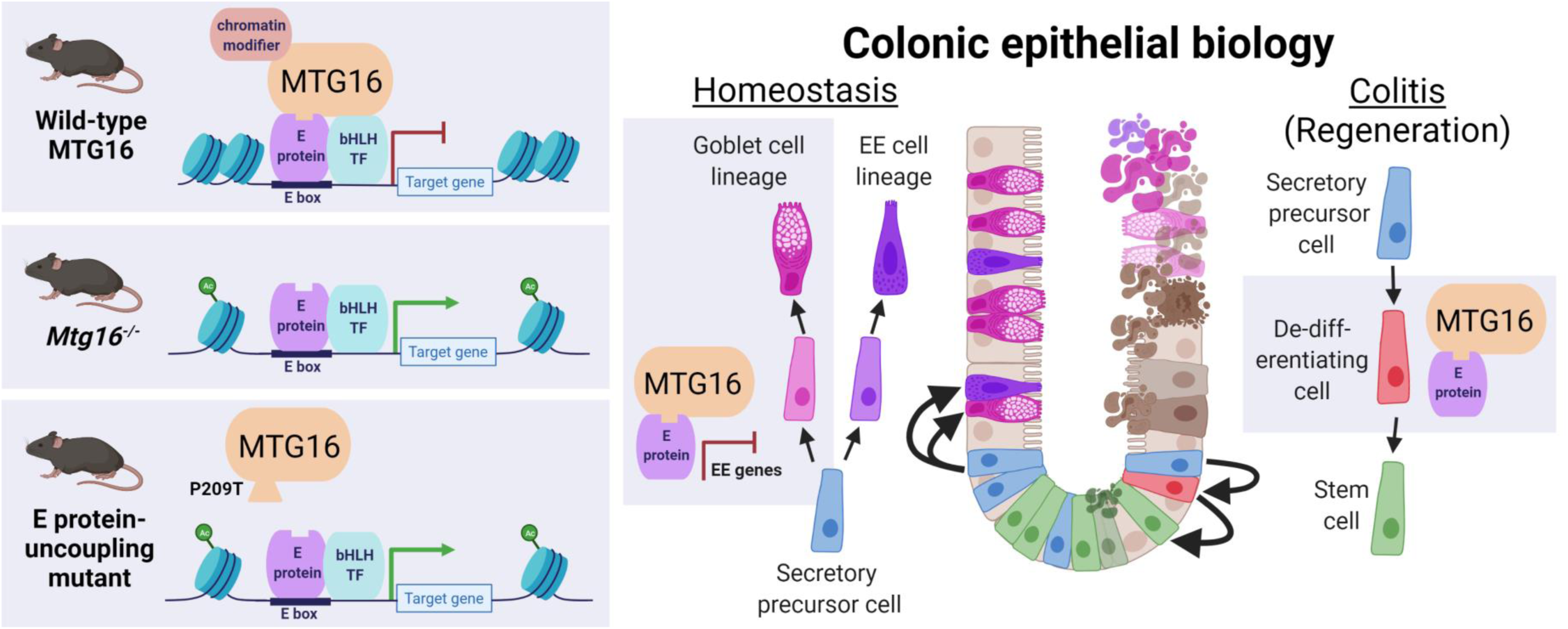

## Introduction

The colonic epithelium is a complex, self-renewing tissue comprised of specialized cell types with diverse functions (1). Stem cells at the base of the colon crypt divide and differentiate into absorptive and secretory cells. Conversely, colonic epithelial regeneration in response to injury occurs through de-differentiation of committed absorptive and secretory lineage cells (2–4). Secretory lineage dysregulation is implicated in inflammatory bowel disease (IBD), diabetes, and even behavioral changes (5–9), and aberrant regenerative programs may lead to uncontrolled cell proliferation and dysplasia (10). Thus, these processes must be tightly controlled by coordinated networks of transcriptional activators and repressors (1).

Myeloid translocation gene 16 (MTG16, also known as CBFA2T3) is a transcriptional corepressor that regulates cell fate decisions by bridging transcription factors and chromatin modifiers to repress transcription of target loci (11–14). Repression targets of MTG16 vary depending on the expression gradients of other components of the repression complex (14, 15). MTG16 represses stem cell genes and pan-secretory genes in the homeostatic small intestine (SI) (16). We have previously shown that MTG16 deficiency leads to increased injury in dextran sulfate sodium (DSS)-induced colitis (17) and tumor burden in azoxymethane (AOM)/DSS-induced inflammatory carcinogenesis and that these were epithelial-specific phenotypes (18). However, the precise mechanisms by which MTG16 controls colonic homeostasis, regeneration, and tumorigenesis remain unknown.

One family of transcription factors repressed by MTG16 are basic helix-loop-helix (bHLH) transcription factors (14, 19–22). Ubiquitously expressed (class I) and tissue-specific (class II) bHLH transcription factors dimerize to control stem cell dynamics, lineage allocation, and differentiation (23–25). Class I bHLH transcription factors are also known as E proteins because they bind to consensus Ephrussi box (E box) DNA sequences and include E12 and E47 (splice variants of *E2A*), HEB, and E2-2 (22, 23). However, the functions of E proteins have mainly been described outside of the colon (26–32).

In this study, we defined the topology of *Mtg16* expression in the colon and discovered new functions of MTG16 in the colonic epithelium. We used a novel MTG16:E protein uncoupling point mutant mouse model to demonstrate that the mechanism driving these phenotypes was repression of E protein transcription factors by MTG16. We tested the functional impact of this regulatory relationship in murine models of IBD and colitis-associated cancer (CAC) and correlated our observations with patient data. Overall, we demonstrate novel, context-specific roles for MTG16 in colonic epithelial lineage allocation and protection from colitis and tumorigenesis that depend on its repression of E protein-mediated transcription.

## Results

### *MTG16* is expressed in goblet cells in human and murine colon

To understand the topography of *MTG16* expression in the human colon, we queried single-cell RNA-seq (scRNA-seq) datasets generated from 2 independent cohorts of normal human colon samples (10). Interestingly, we observed that *MTG16* was predominantly expressed in goblet cells (**Fig. 1A, Fig. S1A**). To understand if this expression pattern was evolutionarily conserved, we queried our inDrop scRNA-seq generated from WT murine colon crypts (33, 34) and similarly found that *Mtg16* was expressed in goblet and enteroendocrine clusters (**Fig. 1B, Fig. S1B**). RNAscope for *Mtg16* and *Muc2*, the predominant mucin expressed by colonic goblet cells (35), demonstrated strong co-localization in the murine colonic epithelium (**Fig. 1C**). Collectively, these data suggested a role for MTG16 in the colonic secretory lineage.

**Figure 1.**
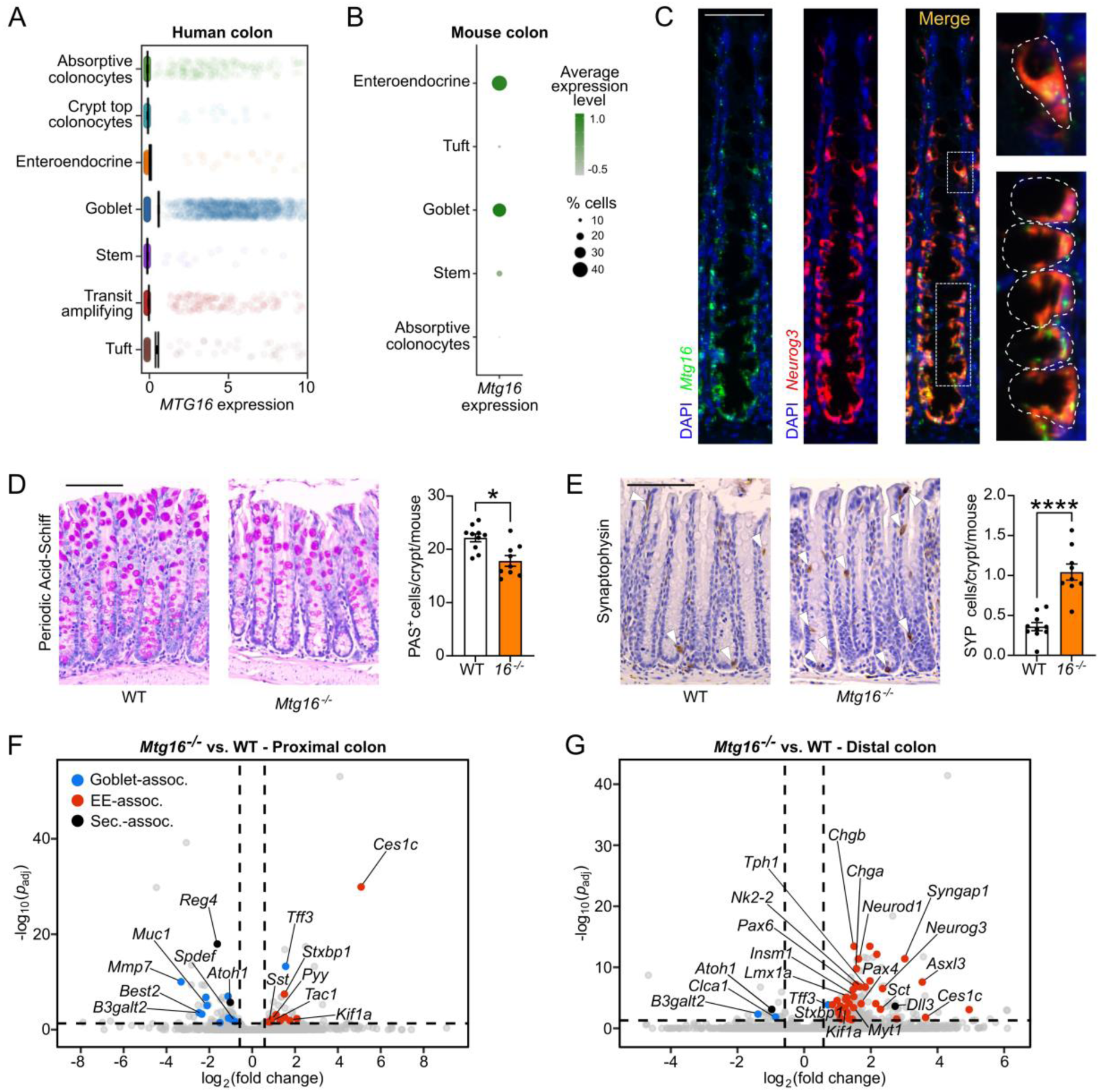
MTG16 is expressed in a subset of colonic secretory cells, and its deficiency alters lineage allocation. **(A)** *MTG16* expression in cell populations generated from inDrop scRNA-seq of normal human colon biopsies (discovery cohort, n = 35 samples, 30,374 cells) (validation cohort in Fig. S1A). **(B)** *Mtg16* expression in inDrop scRNA-seq of WT mouse colon (n = 3 mice, 3,653 cells). Color gradient represents the average *Mtg16* expression level in each cell population. Dot size represents the percentage of cells in each population expressing *Mtg16.* UMAP plots are in Fig. S1B. **(C)** RNAscope *in situ* hybridization of *Mtg16* and *Muc2* in WT mouse colon. Scale bar = 50 µm. Goblet cells are outlined in the insets at right. **(D-E)** WT and *Mtg16^-/-^* mouse colon (n = 10 WT, 9 *Mtg16^-/-^*) stained for **(D)** goblet cells by periodic acid-Schiff (PAS) stain and **(E)** enteroendocrine cells by IHC for synaptophysin (SYP). Representative images at left. SYP^+^ cells are indicated with white arrowheads. Scale bars = 100 µm. \**p* < 0.05, \*\*\*\**p* < 0.0001 by Mann-Whitney test. **(F-G)** Volcano plots demonstrating differentially expressed genes in **(F)** proximal (n = 2 WT, 2 *Mtg16^-/-^*) and **(G)** distal (n = 4 WT, 4 *Mtg16^-/-^*) colonic epithelial isolates by RNA-seq. Horizontal dashed line indicates *p*_adj_ < 0.05 by DESeq2. Vertical dashed lines indicate fold change = 1.5. Dot colors indicate goblet-associated (blue), enteroendocrine (EE)-associated (red), and secretory-associated (black) genes.

### Loss of MTG16 distorts colonic secretory cell differentiation

We next assessed secretory lineage allocation in the homeostatic colon. *Mtg16^-/-^* mice had fewer goblet cells but, interestingly, more synaptophysin-positive (SYP^+^) enteroendocrine cells per colon crypt (**Fig. 1D-E**). Tuft cells were decreased in the colons of *Mtg16^-/-^* mice (**Fig. S1C**). In agreement with prior studies (16, 36), *Mtg16^-/-^* mice had fewer goblet cells, but no increase in enteroendocrine cells, in the SI (**Fig. S2**). Together, these data implicated a colon-specific role for MTG16 in orchestrating cell type allocation within the secretory lineage.

Prior studies have demonstrated subtle differences in secretory cell differentiation along the length of the colon(35, 37–40). Thus, we stratified our human scRNA-seq datasets by proximal and distal colon and examined *MTG16* expression. *MTG16* was consistently expressed in goblet cells, but its expression pattern in other secretory cell types varied (**Fig. S3**). We next queried the *Tabula Muris* (41), a murine colon scRNA-seq dataset that specifies whether certain clusters originate from the proximal or distal colon, for *Mtg16* expression. In the proximal colon, *Mtg16* was enriched in all goblet cell clusters (**Fig. S4A**). In the distal colon, *Mtg16* was enriched in *Lgr5*^+^ amplifying, undifferentiated cells and most goblet cells, except those located at the top of the crypt (**Fig. S4B**). Bulk RNA-seq of proximal and distal WT colon revealed higher *Mtg16* expression in the proximal colon, potentially reflecting its higher goblet cell density (**Fig. S4C**). We confirmed this *Mtg16* expression pattern by RNAscope (**Fig. S5**).

Due to these differences, we performed RNA-seq of colon epithelial isolates from the proximal and distal colon separately. In both colon segments, goblet-associated genes were downregulated, and enteroendocrine-associated genes were upregulated compared to WT (**Fig. 1F-G**). However, goblet cell-associated gene downregulation was more pronounced in the proximal colon (**Fig. 1F**), while enteroendocrine cell-associated gene upregulation was more striking in the distal colon (**Fig. 1G**). Closer inspection of the distal *Mtg16^-/-^* colon revealed marked upregulation of class II bHLH transcription factors required for enteroendocrine cell differentiation (*Neurog3*, *Neurod1*) and enteroendocrine cell hormone products (*Chga, Gcg, Sct*) (**Fig. 2A**). Due to this novel phenotype, we chose to further investigate the distal colon using Gene Set Enrichment Analysis (GSEA) with custom gene sets derived from the literature (described in **Table S3**). GSEA of distal colon RNA-seq demonstrated significant enrichment of all stages of enteroendocrine cells during differentiation except the earliest progenitor cells in which expression of *Neurog3*, which is required for enteroendocrine cell differentiation (42–45), has not yet been induced (44) (**Fig. 2B**). Concurrently, the goblet cell gene signature was significantly de-enriched in the absence of MTG16 (**Fig. 2B**).

**Figure 2.**
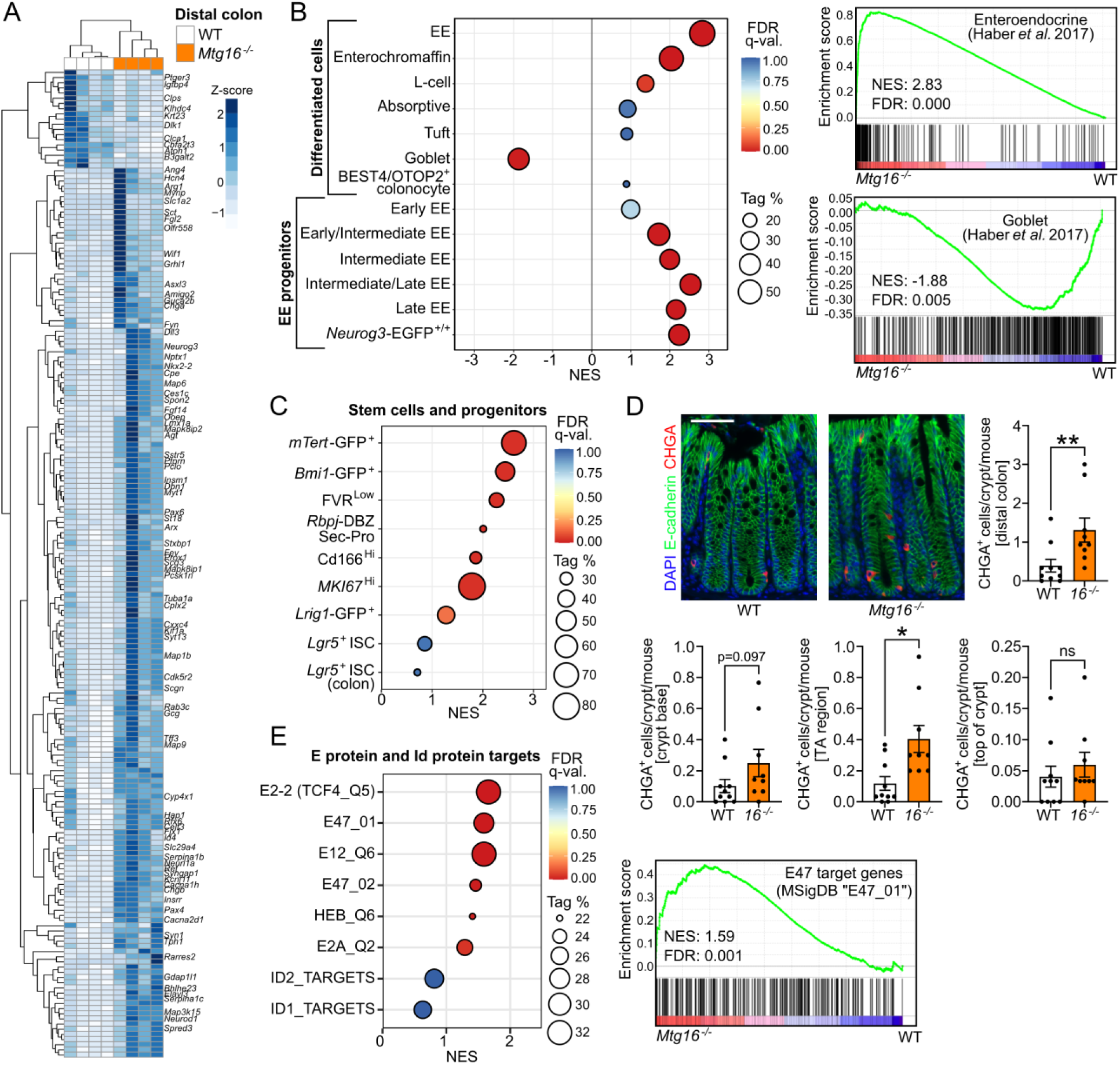
MTG16 deficiency is associated with increased enteroendocrine cell markers, reserve stem cell gene signatures, and E protein target upregulation in the distal colon. **(A)** Unsupervised hierarchical clustering of differentially expressed genes in *Mtg16^-/-^* vs. WT (n = 4) distal colonic epithelial isolates by RNA-seq. Gene expression in individual samples is displayed by Z-score (normalized count). Genes of interest are labeled at right. **(B)** GSEA of distal colon RNA-seq using gene sets representing epithelial cell types (described in Table S3). EE, enteroendocrine. Individual GSEA plots of enteroendocrine and goblet cell signatures are displayed at right. **(C)** GSEA of distal colon RNA-seq using gene sets representing stem and other progenitor cell populations (Table S3). **(D)** Staining and quantification of CHGA^+^ cells/crypt (top) stratified by location along the crypt axis (bottom). Scale bar = 50 µm. n = 10 WT, 9 *Mtg16^-/-^*. \**p* < 0.05, \*\**p* < 0.01 by Mann-Whitney test. **(E)** GSEA of distal colon RNA-seq using MSigDB (95) gene sets representing E protein and Id protein transcriptional signatures (Table S3). Individual GSEA plot of the E47 target gene signature at right. **(B-C, E)** NES, normalized enrichment score (ES). Tag % is defined as the percentage of gene hits before (for positive ES) or after (for negative ES) the peak in the running ES, indicating the percentage of genes contributing to the ES. FDR q-value < 0.05 is considered significant.

Notably, *Mtg16^-/-^* vs. WT RNA-seq did not show significant differences in the stem cell genes *Lgr5* and *Ascl2*, distinct from *Mtg16^-/-^* SI (16). Additionally, GSEA using gene sets representing stem cells and other early progenitors (**Table S3**) did not demonstrate enrichment of any gene sets representing *Lgr5*^+^ crypt base columnar stem cells (CBCs) (**Fig. 2C**, **Fig. S6A**). We did observe enrichment of gene sets representing *Bmi1*^+^ “label-retaining” and *mTert*^+^ stem cells (**Fig. 2C**), recently shown to be enteroendocrine cells expressing chromogranin A (*Chga*) (46). While CHGA^+^ cells were increased overall, they were not localized to the crypt base (including the +4/+5 cell position) (**Fig. 2D**). Altogether, these data indicate that MTG16 promotes goblet cell differentiation, at the expense of enteroendocrine cell differentiation, during colonic secretory lineage allocation.

### MTG16-driven goblet cell differentiation is dependent on its repression of E protein-mediated transcription

We next sought to illuminate potential mechanisms by which MTG16 promotes the goblet lineage and restricts the enteroendocrine lineage. We and others have previously shown that MTG16 negatively regulates major transcription factor effectors of the WNT and Notch pathways in the hematopoietic system (15, 47) and SI (16). We thus performed GSEA for WNT and Notch activation, as well as other pathways important in colonic epithelial homeostasis and differentiation (1). Surprisingly, GSEA did not identify global canonical WNT or Notch pathway upregulation in *Mtg16^-/-^* colon crypts (**Fig. S6B**), despite upregulation of the Notch ligand *Dll3* (**Fig. 1G**, **Fig. 2A**). Additionally, several WNT pathway inhibitors (*Wif1*, *Cxxc4*) and the Notch pathway inhibitor *Neurl1a* were significantly upregulated (**Fig. 2A**). Thus, it appeared unlikely that the effects of MTG16 on colonic differentiation were due to interaction with, and thus repression of, canonical WNT and Notch effectors.

We then investigated whether target genes of known MTG16-interacting factors, including E proteins (14, 19–22), were dysregulated in *Mtg16^-/-^* colon crypts. As stated previously, E proteins coordinate lineage-specific differentiation by interacting with cell type-specific class II bHLH transcription factors (23). Furthermore, E proteins such as E2A and HEB are known to be expressed in SI crypts and adenomas (48). Here, we observed enrichment of all E protein gene signatures by GSEA in *Mtg16^-/-^* colon crypts (**Fig. 2E**, **Table S3**). Gene sets associated with upregulation of Id1 and Id2, which counteract E protein activity and increase stemness in colon cancer (23, 49–52), were not enriched (**Fig. 2E**). These data demonstrated a strong induction of E protein-mediated transcription in the absence of repression by MTG16. Thus, we hypothesized that MTG16 drives goblet cell differentiation from secretory progenitors by inhibiting E protein-mediated transcription of key enteroendocrine transcription factors.

We next tested this hypothesis by selectively disrupting the MTG16:E protein interface *in vivo*. To do so, we leveraged a CRISPR-generated mouse with a point mutation resulting in a proline-to-threonine substitution in MTG16 (*Mtg16^P209T^*) that impairs MTG16:E2A and MTG16:HEB binding and, consequently, MTG16-mediated repression of E protein transcriptional activity (20). Similar to *Mtg16^-/-^* mice, homozygous *Mtg16^P209T^* mutant (*Mtg16^T/T^*) mice had fewer periodic acid-Schiff-positive (PAS^+^) goblet cells and more CHGA^+^ enteroendocrine cells per crypt compared to littermate-matched WT controls, though tuft cell frequency per crypt was unchanged (**Fig. 3A-C**). We next performed bulk RNA-seq on proximal and distal colon crypt isolates. In both colon segments, goblet cell genes were downregulated compared to WT, while enteroendocrine cell genes were upregulated compared to WT, in a pattern resembling that of *Mtg16^-/-^* colon (**Fig. 3D-E**). Notably, the distal colon exhibited significant upregulation of *Neurog3* (**Fig. 3E**). Additionally, GSEA of the overall enteroendocrine signature, key enteroendocrine progenitor genes, *Bmi1*^+^ and *mTert*^+^ stem cell signatures, and E protein signatures all largely phenocopied the *Mtg16^-/-^* distal colon (**Fig. 3F-H, Table S3**). As expected, the gene sets representing E2A and HEB transcriptional targets were also enriched (**Fig. 3H**).

**Figure 3.**
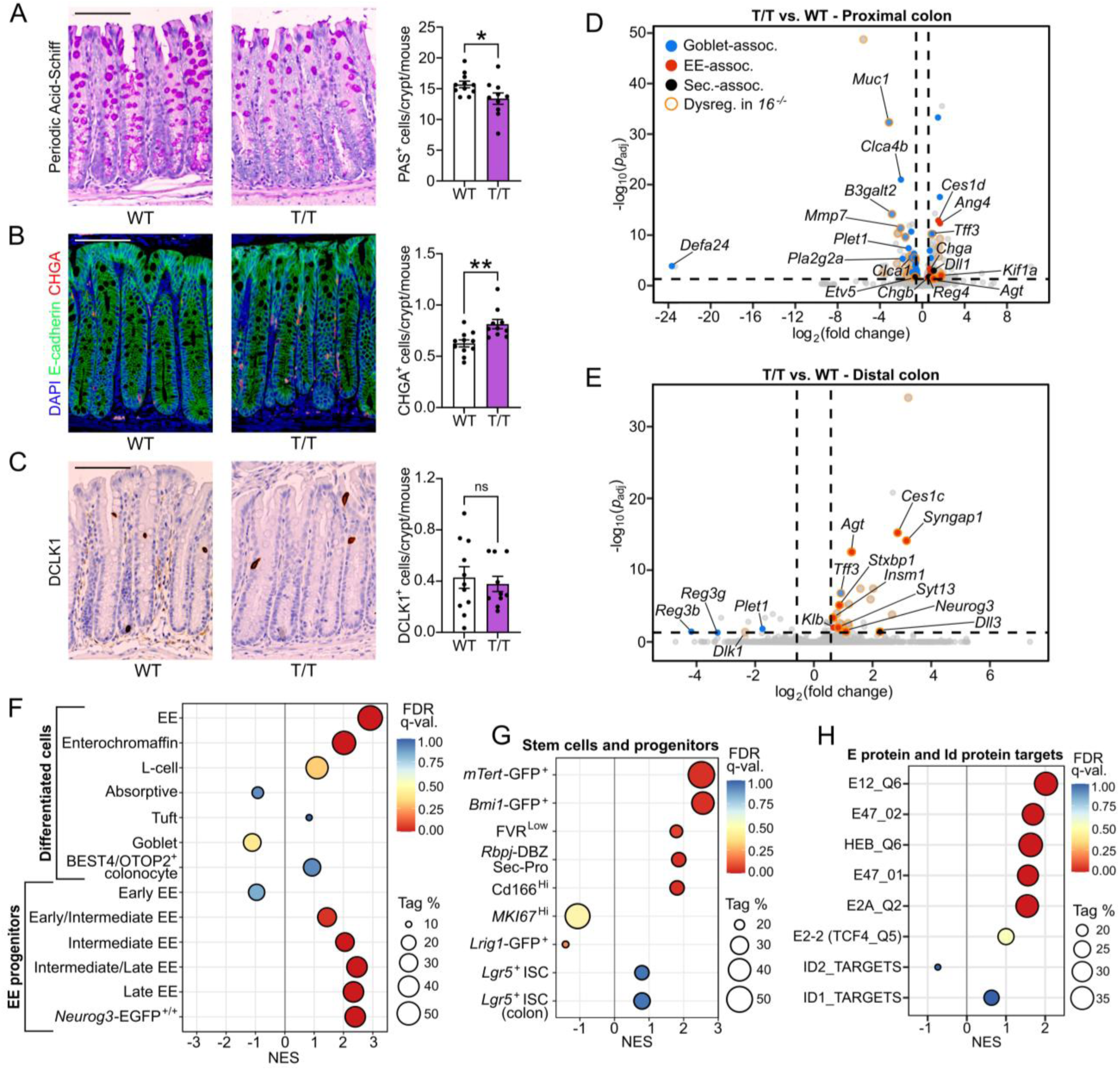
MTG16-driven colonic epithelial lineage allocation is dependent on its repression of the E protein transcription factors E2A and HEB. **(A)** WT and *Mtg16^T/T^* (T/T) mouse colon (n = 11 WT, 10 *Mtg16^T/T^*) stained for **(A)** goblet cells by periodic acid-Schiff (PAS) stain, **(B)** enteroendocrine cells by IHC for synaptophysin (SYP), and **(C)** tuft cells by IHC for doublecortin-like kinase 1 (DCLK1). Representative images at left. Scale bars = 100 µm. \**p* < 0.05, \*\**p* < 0.01 by Mann-Whitney test. **(D-E)** Volcano plots demonstrating differentially expressed genes in **(D)** proximal (n = 2 WT, 2 *Mtg16^T/T^*) and **(E)** distal (n = 3 WT, 4 *Mtg16^T/T^*) colonic epithelial isolates by RNA-seq. Horizontal dashed line indicates *p*_adj_ < 0.05 by DESeq2. Vertical dashed lines indicate fold change = 1.5. Dot colors indicate goblet-associated (blue), enteroendocrine (EE)-associated (red), and secretory-associated (black) genes. Dots outlined in orange represent genes that are upregulated or downregulated in the same direction in the corresponding *Mtg16^-/-^* colon segment (“Dysreg. in *16^-/-^*”). **(F-H)** GSEA of distal colon RNA-seq using gene sets representing **(F-G)** epithelial cell types and **(H)** E protein transcriptional signatures (Table S3). EE, enteroendocrine. NES, normalized enrichment score (ES). Tag % is defined as the percentage of gene hits before (for positive ES) or after (for negative ES) the peak in the running ES, indicating the percentage of genes contributing to the ES. FDR q-value < 0.05 is considered significant.

We next compared RNA-seq results from *Mtg16^-/-^* vs. WT and *Mtg16^T/T^* vs. WT distal colon crypts and observed 22 genes upregulated in both datasets (**Fig. 4A, Table 1**). Many of these genes (e.g. *Neurog3, Insm1, Insrr, Pex5l, Syt13, Syngap1, Kcnh6, and Agt*) were enteroendocrine-associated (**Table 1**). A publicly available MTG16 ChIP-seq dataset from SI crypts (16) exhibited significant MTG16 occupancy proximal to 11 of the 22 genes (**Table 1**), including a large peak at an E box-rich site upstream of *Neurog3* (**Fig. 4B**), suggesting that these genes are repressed by MTG16. Finally, *Neurog3^+^* enteroendocrine progenitor cells were markedly increased in *Mtg16^-/-^* and *Mtg16^T/T^* colon crypts by RNAscope (**Fig. 4C-D**). Altogether, we discovered an MTG16:E protein-dependent mechanism of colonic secretory lineage allocation, specifically inhibition of NEUROG3-mediated commitment to the enteroendocrine cell lineage.

**Figure 4.**
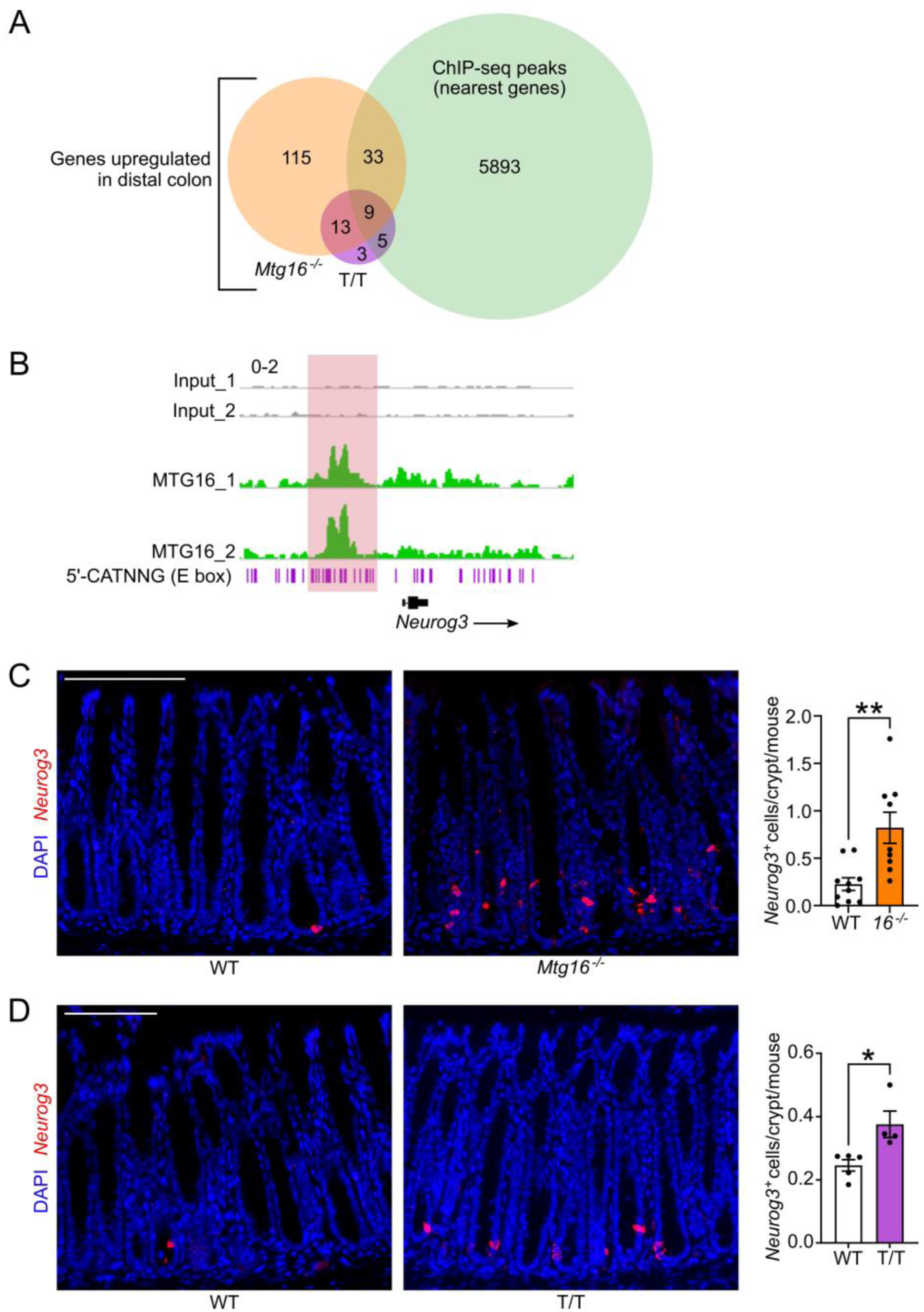
MTG16 represses E protein-mediated transcription of *Neurog3*. **(A)** Venn diagram of upregulated (*p*_adj_ < 0.05) genes from bulk RNA-seq of WT vs. *Mtg16^-/-^* (orange) and WT vs *Mtg16^T/T^* (T/T) (purple) distal colonic crypt isolates (n = 3-4) and genes with transcription start sites (TSS) nearest to MTG16 peaks in a ChIP-seq dataset generated by Baulies *et al.* (16) (green). Table 1 provides more information about the common upregulated genes. **(B)** MTG16 occupancy of an E box (5’-CATNNG)-rich region 5’ to *Neurog3* in ChIP-seq by Baulies *et al.* (16). Scale: 0-2. **(C-D)** Staining and quantification of *Neurog3*^+^ enteroendocrine progenitor cells by RNAscope in **(C)** WT vs. *Mtg16^-/-^* colon (n = 9-10) and **(D)** WT vs. *Mtg16^T/T^* colon (n = 4-5). \**p* < 0.05, \*\**p* < 0.01 by Mann-Whitney test.

**Table 1.** Genes upregulated in both *Mtg16^-/-^* vs. WT and *Mtg16^T/T^* vs. WT distal colon RNA-seq. Log2FC. The log_2_(fold change) (log_2_FC) of each gene in each dataset (*Mtg16^-/-^* vs. WT distal colon and *Mtg16^T/T^* vs. WT distal colon) and whether they have a TSS nearest to an MTG16 ChIP-seq peak in a dataset generated by Baulies et al. (16) is shown.

| Genes up in<br>distal colon | *Mtg16*<sup>-/-</sup> vs. WT<br>(log<sub>2</sub>FC) | *Mtg16*<sup>T/T</sup> vs. WT<br>(log<sub>2</sub>FC) | ChIP-seq<br>peak<br>(nearest<br>TSS)? |
| --- | --- | --- | --- |
| *Agt* | 1.55 | 1.29 | No |
| *Ces1c* | 2.77 | 2.85 | No |
| *Dbn1* | 2.07 | 0.95 | No |
| *Dll3* | 2.73 | 2.25 | Yes |
| *Dmpk* | 1.80 | 1.26 | Yes |
| *Gfra3* | 1.07 | 0.89 | No |
| *Gnao1* | 1.42 | 1.20 | No |
| *Guca2b* | 1.14 | 0.79 | Yes |
| *Id4* | 1.92 | 1.19 | No |
| *Insm1* | 1.49 | 0.65 | No |
| *Insrr* | 4.30 | 3.22 | Yes |
| *Kcnh6* | 1.94 | 1.17 | Yes |
| ***Neurog3*** | **2.35** | **1.09** | **Yes** |
| *Ooep* | 2.83 | 2.66 | No |
| *Pex5l* | 2.65 | 2.02 | No |
| *Prss23* | 2.52 | 1.59 | Yes |
| *Rbp4* | 1.66 | 0.88 | No |
| *Spon2* | 2.84 | 1.93 | No |
| *Stxbp1* | 1.04 | 0.87 | Yes |
| *Syngap1* | 3.02 | 3.16 | No |
| *Syt13* | 1.97 | 0.83 | No |
| *Tff3* | 0.72 | 0.92 | Yes |

### *MTG16* is upregulated in IBD patients with active disease

There has been considerable interest in the role of secretory cell lineage misallocation and dysfunction in the pathogenesis of IBD, which includes ulcerative colitis (UC) and Crohn’s disease and is defined by continuous epithelial injury and regeneration (8, 35, 53–57). Additionally, several large RNA-seq studies of human UC biopsies have recently been published, including the pediatric PROTECT study (58, 59) and mucosal biopsies from adult UC patients with active disease, UC patients in remission, and healthy control patients (60). We found that *MTG16* was upregulated in UC patients in both datasets (**Fig. 5A-B**). *MTG16* expression decreased with remission, although remained elevated compared to unaffected control patients (**Fig. 5B**). Lastly, a microarray dataset from pediatric colon biopsies (61) demonstrated upregulation of *MTG16* in both UC and Crohn’s colitis (**Fig. 5C**). Collectively, these data show that increased *MTG16* expression is correlated with active injury and regeneration in IBD.

**Figure 5.**
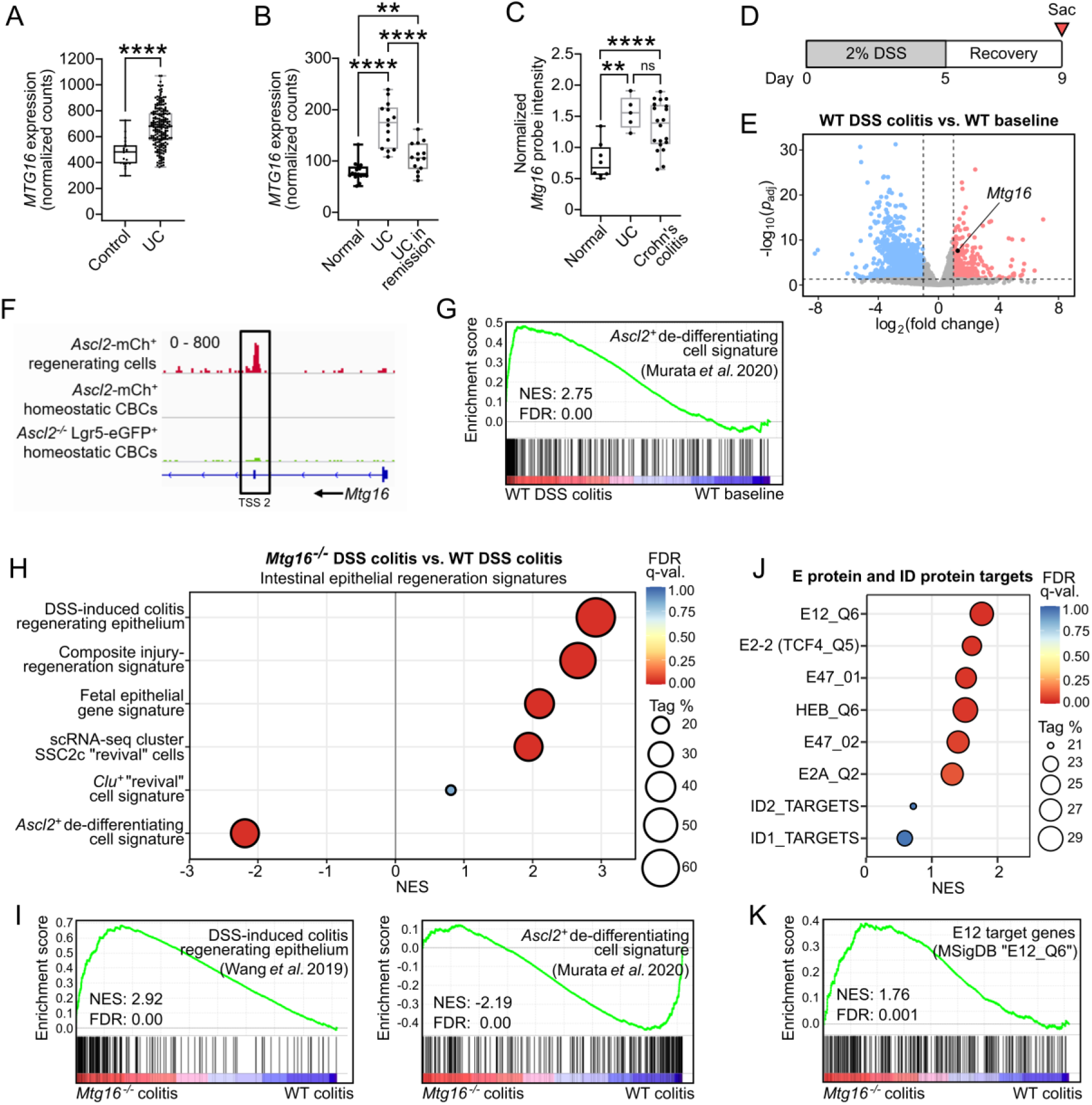
*Mtg16* is upregulated in experimental colitis and patients with IBD and is required for ASCL2-mediated colonic epithelial regeneration. **(A)** *MTG16* expression from an RNA-seq dataset generated from rectal biopsies of treatment-naïve pediatric UC patients (n = 206) vs. unaffected controls (n = 20) in the PROTECT study (58). Transcript counts were normalized by DESeq2. \*\*\*\**p*_adj_ < 0.0001 by DESeq2. **(B)** *MTG16* expression in an RNA-seq dataset from adult ulcerative colitis (UC) patients with active disease (n = 14), UC patients in remission (n = 14), and unaffected control patients (n = 16) (60). Transcript counts were normalized by DESeq2. \*\**p*_adj_ < 0.01 and \*\*\*\**p*_adj_ < 0.0001 by pairwise DESeq2. **(C)** *MTG16* expression in a microarray dataset comparing UC (n = 5), Crohn’s colitis (n = 20), and unaffected (n = 8) patient biopsies (61). \*\**p*_adj_ < 0.01 and \*\*\*\**p*_adj_ < 0.0001 by limma in GEO2R. Normalized probe intensity is plotted for data visualization. **(A-C)** The lines within each box represent the mean, the bounds of the boxes represent 25^th^ to 75^th^ %-iles, and the bounds of the whiskers represent the range of the data. All data points are shown. **(D)** Schematic of the dextran sulfate sodium (DSS) injury-regeneration model in which mice are treated with 2% for 5 d followed by 4 d of recovery. **(E)** Volcano plot demonstrating significantly downregulated (blue) and upregulated (red) genes in distal colon crypt isolates from WT mice following DSS-induced injury and regeneration compared to baseline (n = 3). Horizontal dashed line indicates *p*_adj_ < 0.05 by DESeq2. Vertical dashed lines indicate fold change = 2. **(F)** ASCL2 occupancy near an *Mtg16* transcription start site (TSS 2) in a ChIP-seq dataset of FACS-sorted *Ascl2*^+^ regenerating cells compared to *Lgr5^+^* homeostatic crypt-base columnar stem cells (CBCs) generated by Murata *et al.* (2). Scale: 0-800. **(G)** GSEA plot demonstrating significant enrichment of the *Ascl2*^+^ de-differentiating cell signature (Table S3) in in distal colon crypt isolates from WT mice following DSS-induced colitis vs. WT mice at baseline (n = 3). **(H-K)** GSEA performed on RNA-seq of *Mtg16^-/-^* vs. WT distal colon crypt isolates following DSS injury-regeneration (n = 3 WT, 4 *Mtg16^-/-^*) using **(H-I)** epithelial regeneration-associated gene sets and **(J-K)** gene sets representing E protein transcriptional signatures (described in Table S3). **(G-K)** NES, normalized enrichment score (ES). Tag % is defined as the percentage of gene hits before (for positive ES) or after (for negative ES) the peak in the running ES, indicating the percentage of genes contributing to the ES. FDR q-value < 0.05 is considered significant.

### MTG16 contributes to colon crypt regeneration following DSS-induced colitis

As stated previously, MTG16 deficiency increases disease severity in DSS-induced colitis (17), a mouse model of IBD (schematic in **Fig. 5D**) that recapitulates histologic features of human UC (62). We first assessed *Mtg16* expression in DSS-induced colitis to determine whether it would recapitulate the increased *MTG16* expression in human IBD. Because DSS treatment largely affects the distal colon (62), we compared distal WT colon crypt epithelial isolates post-DSS-induced injury and regeneration to WT distal colon crypts at baseline. Indeed, *Mtg16* expression was significantly increased in regenerating colon crypts (**Fig. 5E**), similar to human IBD. Thus, we posited that MTG16 expression may be critical for epithelial regeneration in colitis.

The cellular sources of intestinal repair have been a major focus of the field over the last several decades. Recently, Murata *et al.* demonstrated that any CBC progeny cell can de-differentiate and repopulate ablated CBCs by inducing expression of *Ascl2* (2). After CBC ablation, 238 significantly upregulated genes defined the de-differentiating ASCL2^+^ cells compared to resting CBCs. Interestingly, *Mtg16* was one of these few significantly upregulated genes (log_2_[fold change] = 2.05, *p*_adj_ = 0.004) (2). Furthermore, ASCL2 ChIP-seq data in the de-differentiating population (2) yielded a peak at a second transcription start site of *Mtg16*, suggesting that ASCL2 induces *Mtg16* expression during colonic epithelial regeneration (**Fig. 5F**).

To confirm that the ASCL2-driven regeneration program was relevant to crypt regeneration following DSS-induced injury, we constructed a gene set consisting of the 238 genes upregulated in *Ascl2*^+^ regenerating cells (2) (**Table S3**), performed GSEA, and observed significant enrichment of this signature in WT distal colon crypts after DSS-induced regeneration (**Fig. 5G**). Conversely, this regeneration signature was significantly de-enriched in *Mtg16^-/-^* colon crypts following DSS-induced injury, despite enrichment of other gene signatures associated with intestinal epithelial regeneration (**Fig. 5H-I**, **Table S3**). Altogether, these data implicate MTG16 as a central component of ASCL2-driven regeneration and repair in the colon. Similar to our findings in colon homeostasis, E protein transcriptional signatures were enriched in post-DSS *Mtg16^-/-^* colon by GSEA (**Fig. 5J-K**), leading us to hypothesize that the contribution of MTG16 to colon crypt regeneration occurs via repression of E protein-mediated transcription.

### MTG16-mediated colonic epithelial regeneration is dependent on its repression of E proteins

To test whether MTG16-dependent regeneration occurs via an E protein-dependent mechanism, we next analyzed colonic epithelial regeneration following DSS-induced injury (**Fig. 6A**) in *Mtg16^T/T^* vs. WT mice. Like *Mtg16^-/-^* mice, *Mtg16^T/T^* mice developed worse colitis and poor regeneration measured by increased weight loss (**Fig. 6B**), decreased colon length (**Fig. 6C**), and increased histologic injury-regeneration score (**Fig. 6D, Table S2**) compared to WT. *Mtg16^T/T^* mice also exhibited increased extent of injury along the length of the *Mtg16^T/T^* colon (**Fig. 6E**). Next, we injected mice with 5-Ethynyl-2’-deoxyuridine (EdU) 1 h prior to sacrifice following DSS injury and regeneration and found that *Mtg16*^T/T^ mice had fewer S-phase cells in injury-adjacent crypts (**Fig. 6F**), indicating a proliferation defect in the crypts that actively regenerate the ulcerated colonic epithelium following DSS-induced colitis (63). Finally, like *Mtg16^-/-^* mice, RNA-seq of *Mtg16^T/T^* distal colon crypt epithelial isolates following DSS injury and regeneration was de-enriched for ASCL2-mediated de-differentiation despite enrichment of other gene signatures associated with regeneration (**Fig. 6G-H, Table S3**). These data showed that MTG16 repression of E protein-mediated transcription is required for ASCL2-mediated colonic epithelial regeneration.

**Figure 6.**
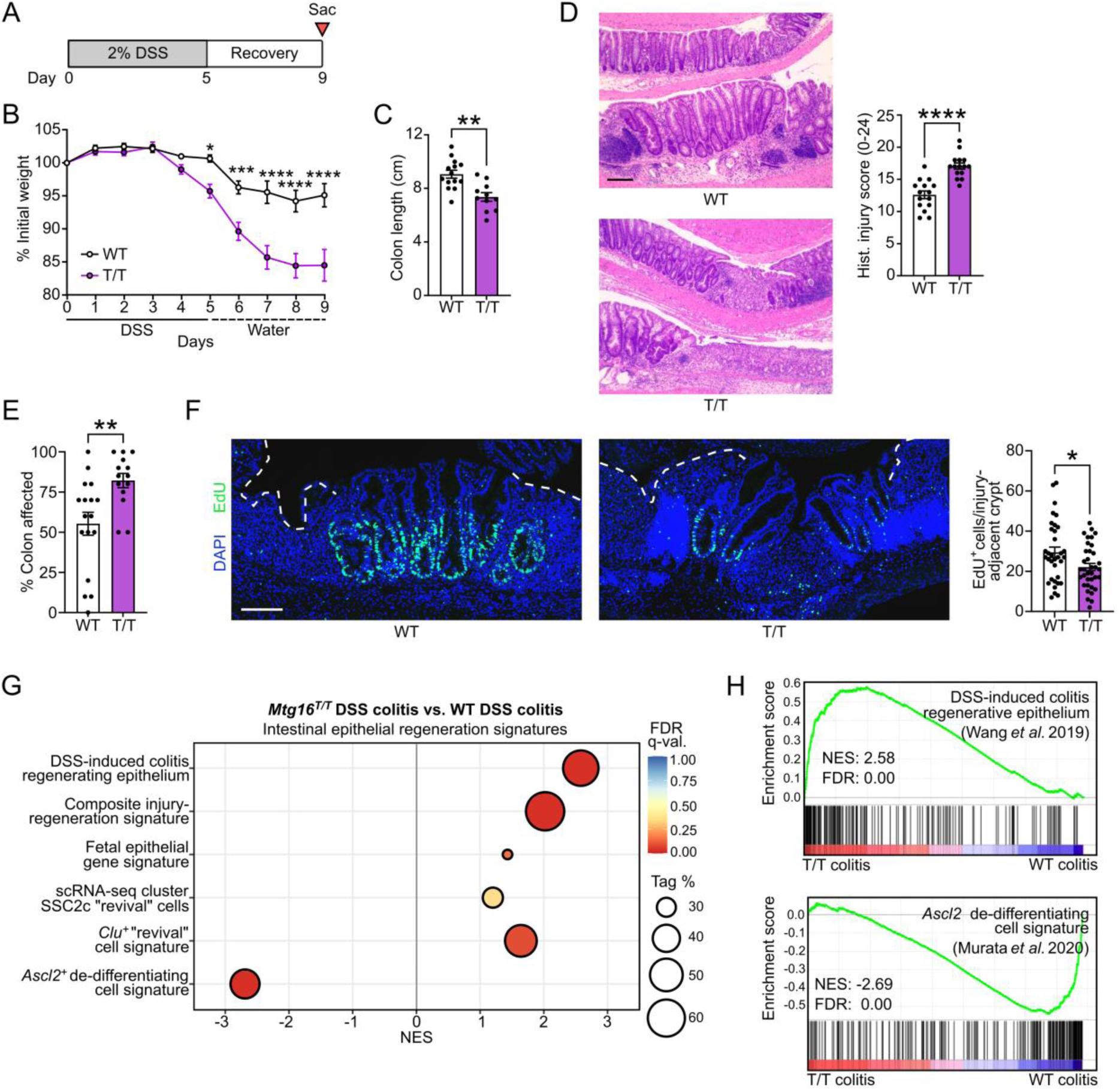
MTG16-mediated colonic epithelial regeneration is dependent on its repression of E protein transcription factor-mediated transcription. **(A)** Schematic of the dextran sulfate sodium (DSS) injury-regeneration model. **(B-D)** Colitis severity in *Mtg16^T/T^* (T/T) vs. WT mice evaluated using **(B)** weight loss, **(C)** colon length, and **(D)** histologic injury (scoring system described in Table S2). **(D)** Representative images at left. Scale bar = 200 µm. **(E)** Percent of colon with histologic findings, estimated from the distal end, as a measure of proximal extent of disease. **(B-E)** Data are combined from 2 independent experiments (n = 15 WT, 16 *Mtg16^T/T^*). **(F)** EdU^+^ cells in injury-adjacent, regenerating crypts in colons from mice injected with EdU 1 h prior to sacrifice on day 9 (n = 30-50 crypts from n = 5 mice). Dashed white line indicates ulcerated epithelium. Scale bar = 100 µm. **(B-F)** \**p <* 0.05, \*\**p <* 0.01, \*\*\**p <* 0.001, \*\*\*\**p <* 0.0001 by **(B)** 2-way ANOVA followed by Sidak’s multiple comparison tests or **(C-F)** Mann-Whitney test. **(G-H)** GSEA of distal *Mtg16^T/T^* vs. WT colon crypt isolates following DSS injury-regeneration (n = 3) (gene sets described in Table S3). NES, normalized enrichment score (ES). Tag % is defined as the percentage of gene hits before (for positive ES) or after (for negative ES) the peak in the running ES, indicating the percentage of genes contributing to the ES. FDR q-value < 0.05 is considered significant.

### *MTG16* expression is decreased in dysplasia

Because *MTG16* expression was upregulated in IBD, we wondered whether *MTG16* expression would be increased in IBD-driven colon cancer (CAC) compared to sporadic CRC. We previously observed decreased *MTG16* expression in sporadic CRC and CAC compared to normal colon tissue (18), but had not directly compared expression levels of *MTG16* between these different neoplastic origins. To do so, we first confirmed our previous findings in sporadic CRC using DESeq2 on raw counts generated by Rahman *et al.* (64) from The Cancer Genome Atlas (TCGA) (65) (**Fig. 7A**). We then analyzed *MTG16* expression in bulk RNA-seq of CRC and CAC tumors (66) and found no significant difference in *MTG16* expression (**Fig. 7B**), indicating that *MTG16* expression is reduced in colon dysplasia regardless of whether it is driven by IBD.

**Figure 7.**
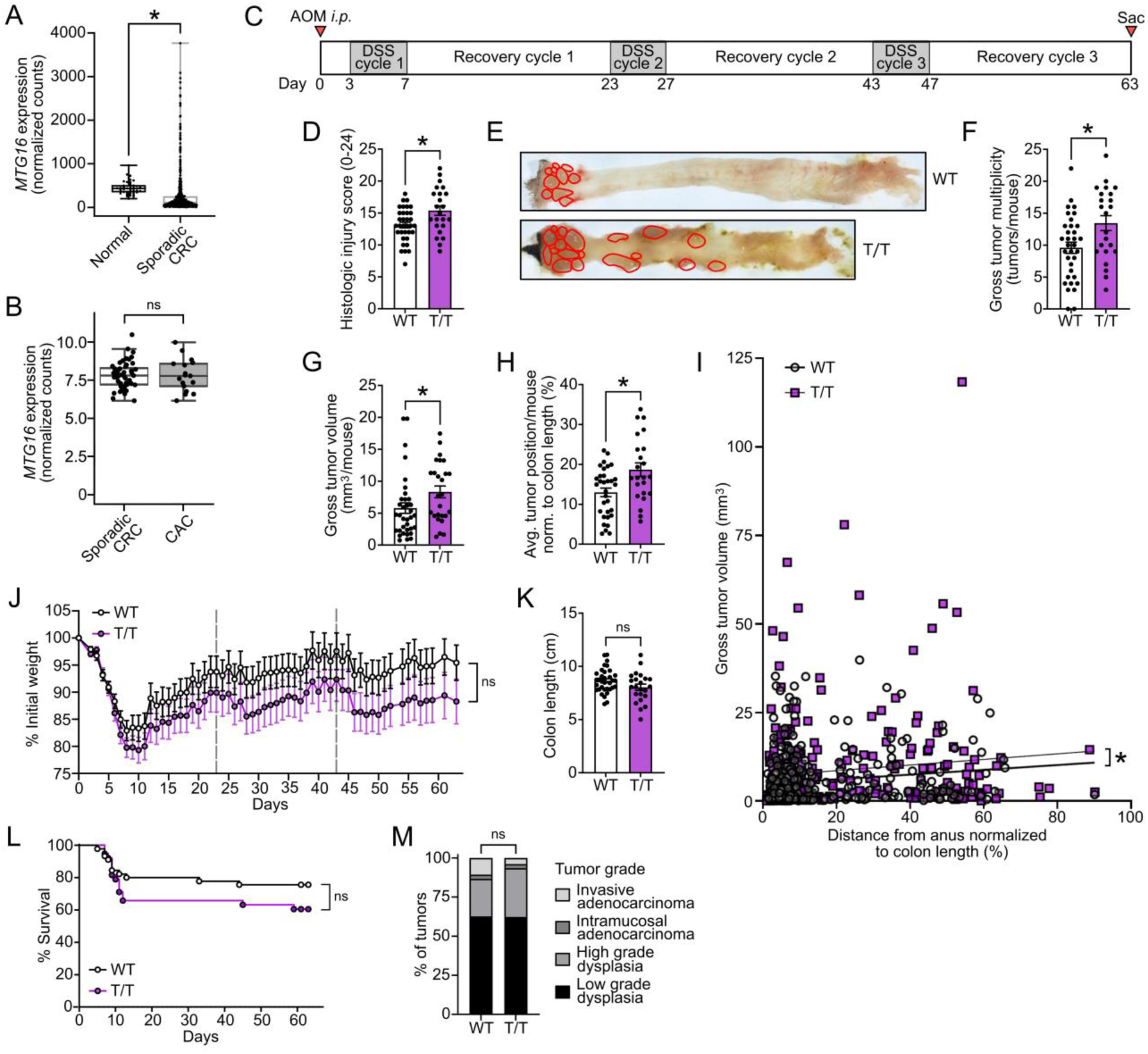
MTG16-mediated protection from tumorigenesis is partially dependent on repression of E protein activity. **(A)** *MTG16* expression in sporadic colorectal cancer (CRC) (n = 482) compared to normal colon tissue (n = 41) by DESeq2 of TCGA raw counts generated by Rahman *et al.* (64). The lines within each box represent the mean, the bounds of the boxes represent the 25^th^ to 75^th^ %-iles, and the bounds of the whiskers represent the range of the data. All individual data points are shown. \**p*_adj_ < 0.05 by DESeq2. **(B)** *MTG16* expression in sporadic CRC (n = 47) and colitis-associated cancer (CAC) (n = 17) tumors. *MTG16* expression was normalized by DESeq2 and limma batch correction. The box extends from the lower to upper quartile values of the data, with a line at the median. The lower whisker is at the lowest datum above Q1 - 1.5*(Q3-Q1), and the upper whisker at the highest datum below Q3 + 1.5*(Q3-Q1), where Q1 and Q3 are the first and third quartiles. **(C)** Schematic of the azoxymethane/dextran sulfate sodium (AOM/DSS) model in which mice are injected *i.p.* with AOM followed by 3 cycles of DSS-induced injury and recovery prior to sacrifice. **(D-G)** CAC severity in AOM/DSS-treated *Mtg16^T/T^* (T/T) vs. WT mice assessed by **(D)** histologic injury-regeneration score (scoring system described in Table S2), **(E-F)** tumor multiplicity, and **(G)** average tumor volume per mouse. **(H)** Average tumor position normalized to colon length (distance from anus/colon length*100%) per mouse. **(J)** Weight loss compared using repeated measures 2-way ANOVA (n = 26 WT, 25 *Mtg16^T/T^*, representative of 2 independent experiments). **(K)** Colon length measured at sacrifice. **(L)** Survival curves compared using Log-rank (Mantel-Cox) test. **(O)** Percent of tumors in each grade of dysplasia, evaluated by a pathologist blinded to genotype and compared by Χ^2^ test. **(D-F, G-I, K-M)** Data are pooled from 2 independent experiments with n = 35 WT, 26 *Mtg16^T/T^* remaining at sacrifice. \**p <* 0.05 by **(D, F-H, K)** Mann-Whitney test or **(I)** least-squares regression.

### MTG16-mediated protection from CAC is partially dependent on its repression of E protein transcription factors

We previously showed that *Mtg16^-/-^* mice have increased mortality, tumor burden, tumor grade, and tumor cell proliferation and apoptosis in AOM/DSS-induced CAC (18). We next tested whether the mechanism driving these phenotypes was E protein-dependent by treating *Mtg16^T/T^* mice with AOM/DSS (**Fig. 7C**). Llike *Mtg16^-/-^* mice, *Mtg16^T/T^* mice did exhibit increased histologic injury, tumor multiplicity, and tumor size compared to WT (**Fig. 7D-G**). Interestingly, *Mtg16^T/T^* mice had more tumors and larger tumors in the middle and proximal colon than their WT counterparts (**Fig. 7L-K**). This was consistent with a greater extent of DSS-induced injury (**Fig. 6E**). These data suggest that this phenotype could be driven by increased injury in the proximal colon.

However, unlike *Mtg16^-/-^* mice, *Mtg16^T/T^* mice did not have significantly more weight loss (**Fig. 7J**), colon shortening (**Fig. 7K**), mortality (**Fig. 7L**), or tumor dysplasia (**Fig. 7M**) compared to WT. Additionally, further characterization of *Mtg16^T/T^* tumors (**Fig. S7**) demonstrated no difference in proliferation (**Fig. S7A**). We did, however, observe a subtle increase in apoptotic epithelial cells per tumor (**Fig. S7B**), similar to our prior findings in *Mtg16^-/-^* tumors. These phenotypes were recapitulated in tumoroids derived from distal, but not proximal, WT and *Mtg16^T/T^* tumors (**Fig. S8**). Overall, these data indicate that uncoupling MTG16:E protein complexes recapitulates some, but not all, *Mtg16^-/-^* phenotypes, suggesting that MTG16 protects from tumorigenesis through additional mechanisms.

## Discussion

In this study, we investigated the mechanism underlying context-specific functions of the transcriptional corepressor MTG16. We identify previously unappreciated roles for MTG16 as a critical regulator of colonic secretory cell differentiation and active colon regeneration. To define the mechanism underlying these phenotypes, we utilized a novel mouse model with a single point mutation in MTG16, *Mtg16^P209T^*, that disrupts MTG16 binding to the E proteins E2A and HEB(20). *Mtg16^T/T^* mice homozygous for this MTG16:E protein uncoupling mutation largely recapitulated *Mtg16^-/-^* lineage allocation and regeneration phenotypes, although the effect of MTG16 loss in dysplasia appears to only partially be due to increased E protein activity. Thus, these data indicate a new role for MTG16:E protein complexes in the colonic epithelium.

First, we demonstrated a novel role for MTG16 in colonic epithelial homeostasis. MTG16 has been shown to repress stem, goblet, and enteroendocrine cell genes to drive enterocyte differentiation in the SI (16). However, in accordance with numerous studies demonstrating important differences between SI and colon biology (2, 35, 67–70), *Mtg16^-/-^* colon did not exhibit increased stem cell transcriptional signatures or an overall decrease in secretory genes. Instead, we observed specific upregulation of the enteroendocrine lineage, occurring at least in part due to unrepressed expression of *Neurog3*, the class II bHLH transcription factor required for enteroendocrine cell differentiation (42–45). This could be due to the fact that MTG16 repression targets, and its effects on them, vary in different tissues and cell types due to expression gradients of its binding partners (14). Another potential explanation for the difference between SI and colonic phenotypes is that *Neurog3^+^* progenitor cells in the SI may still differentiate into goblet and Paneth cells but are fully committed to the enteroendocrine lineage in the colon (45, 68).

Enteroendocrine cell lineage commitment is dependent on a cascade of type II bHLH transcription factors including NEUROG3 and its downstream target NEUROD1 (71). E proteins are known to induce transcription of *Neurog1-3* in neurogenesis (30, 32), and NEUROD1 heterodimerizes with E proteins in other systems (72). Although E proteins have been detected in SI stem cells and adenomas (48), E protein function in colon homeostasis remained unknown. Here, we mechanistically probed the extent to which MTG16:E protein interactions drive MTG16-associated lineage allocation functions in the colon using a novel mouse model with a single amino acid substitution in *Mtg16* (*Mtg16*^P209T^) (20). Strikingly, *Mtg16^T/T^* mice recapitulated secretory cell phenotypes of *Mtg16^-/-^* mice including decreased goblet cells, increased enteroendocrine cells, and increased *Neurog3* tone. One potential non-E protein-mediated pathway that was increased (FDR q-value = 0.11) in *Mtg16^-/-^*, but not *Mtg16^T/T^* colon crypts, is the non-canonical, β-catenin-independent WNT-planar cell polarity (PCP) pathway. This could be responsible for the slight difference in magnitude between *Mtg16^-/-^* and *Mtg16^T/T^* phenotypes, as the WNT-PCP pathway may enable enteroendocrine cells to differentiate directly from “unipotentially-primed” intestinal stem cells^99^.

An area of considerable interest in gastrointestinal biology is the relationship between epithelial differentiation and regeneration (4). Although it was long thought that dedicated *Bmi1*^+^ or *mTert*^+^ “reserve” stem cells at the +4/+5 position replenished the stem cell compartment in intestinal epithelial regeneration, recent data indicate that these cells are enteroendocrine cells that can de-differentiate into stem cells following injury (46). Recently, “committed” progenitors from both the secretory and absorptive lineages were demonstrated to share this ability through induction of ASCL2 (2). In this study, *Ascl2*^+^ de-differentiating cells were isolated from the colonic epithelium following CBC ablation (2). Interestingly, these cells were not enriched for *Clu* or fetal epithelial genes (2), previously thought to enhance intestinal epithelial response to injury (4, 73, 74). *Mtg16* was one of the few genes upregulated in these regenerating cells, and ChIP-seq demonstrated an ASCL2 peak at a second TSS in the *Mtg16* locus (2). Demonstrating relevance to DSS-induced colon regeneration, both *Mtg16* and the *Ascl2^+^* regenerating cell signature were upregulated in regenerating WT colon epithelium compared to baseline. *MTG16* was also upregulated in multiple IBD patient cohorts and even decreased with remission, during which the colon epithelium is closer to homeostatic renewal. Finally, RNA-seq and GSEA of *Mtg16^-/-^* and *Mtg16^T/T^* colonic epithelial isolates following DSS-induced injury demonstrated enrichment of “revival” stem cell signatures and fetal epithelial signatures, but de-enrichment of the *Ascl2^+^* regenerating cell signature. These data indicate an active, E protein-mediated role for MTG16 in colon regeneration and provide a potential explanation for defective colonic epithelial regeneration in *Mtg16^-/-^* mice despite an apparent baseline enrichment of *Bmi1*^+^ and *mTert*^+^ “reserve” stem cells. Future work is necessary to determine the E protein targets repressed by MTG16 in *Ascl2^+^* de-differentiating cells. One possibility is that MTG16 represses *Neurog3* in these de-differentiating cells to prevent them from reversing course and differentiating toward the enteroendocrine lineage.

Finally, we investigated the role of MTG16:E protein complexes in inflammatory carcinogenesis. We previously showed that *Mtg16^-/-^* mice develop greater tumor burden than their WT counterparts with AOM/DSS treatment (18). Although E proteins maintain lineage commitment in homeostasis, they have been shown to be both anti-tumorigenic and pro-tumorigenic in colon cancer (75, 76). Although we did observe increased histologic injury, tumor burden, and tumor size in *Mtg16^T/T^* mice, *Mtg16^T/T^* tumors did not recapitulate the much higher mortality or grade of dysplasia observed in *Mtg16^-/-^* tumors (18). Thus, disabling one known function of MTG16 was not sufficient to fully replicate the *Mtg16^-/-^* phenotype. Indeed, our group recently showed that MTG16-dependent effects on CAC development is dependent on Kaiso (ZBTB33) (77). Future studies are necessary to determine additional MTG16 binding partners and repression targets responsible for suppressing carcinogenesis.

In conclusion, we discovered key new functions of MTG16 in colonic secretory lineage allocation, regeneration following colitis, and colitis-associated tumorigenesis, dependent on its repression of E proteins. Confirming translational relevance, we determined that *MTG16* is upregulated in patients with active IBD, reduced with restitution, and decreased in dysplasia. Thus, *MTG16* may be a candidate biomarker for disease activity or a target to modulate differentiation and regeneration.

## Methods

### Animal models

*Mtg16^-/-^* mice were previously generated and validated (78). *Mtg16^P209T^* mutant mice were CRISPR-generated as recently described (20). Both models are on a predominantly C57BL/6 background. Littermate-matched WT and *Mtg16^-/-^* and WT and homozygous *Mtg16^P209T^* mutant (*Mtg16^T/T^*) mice were bred using heterozygous x heterozygous breeding schemes, housed in the same facility with a 12:12 light-dark cycle, provided with standard rodent chow (#5L0D, LabDiet) *ad libitum*. Approximately equal numbers of 8-12-wk-old male and female mice were used.

### scRNA-seq

InDrop scRNA-seq and analysis were performed on 2 independent (DIS, discovery and VAL, validation) cohorts of human colon samples as previously described (10). These data are publicly available through the Human Tumor Atlas Network (HTAN) (https://data.humantumoratlas.org/) (79). scRNA-seq of colon crypts isolated from WT mice was performed as previously described (33, 34, 80). These data (#GSE145830, #GSE114044) are available in the Gene Expression Omnibus (GEO) (81, 82).

### RNAscope

RNAscope, high-resolution RNA *in situ* hybridization (83), was performed on FFPE sections using the RNAscope Multiplex Fluorescent V2 Assay (#323100, ACDBio). Slides were boiled in Target Retrieval Reagents for 15 min and treated with Protease IV for 30 min. Probes were specific for Mm-*Cbfa2t3* (#43601-C2, ACDBio), Mm-*Muc2* (#315451, ACDBio), and Mm-*Neurog3* (#422401, ACDBio). 1:750 TSA Cy3 (#NEL704A001KT, PerkinElmer) and TSA Cy5 (#NEL75A001KT, Perkin Elmer) were used for probe visualization.

### Chromogenic and immunofluorescent IHC

SI and colon were dissected, Swiss-rolled, and fixed in 10% neutral-buffered formalin (NBF) for 24 h at RT. formalin (NBF) for 24 h at RT. The Vanderbilt Translational Pathology Shared Resource (TPSR) paraffin-embedded the tissue and performed H&E and PAS staining. Chromogenic and immunofluorescent IHC were performed as previously described (77, 84). Briefly, 5-µm FFPE sections were deparaffinized, rehydrated in graded ethanol concentrations, and permeabilized using TBS-T (Tris-buffered saline with 0.05% Tween-20) followed by antibody-specific antigen retrieval (**Table S1**). Next, for chromogenic IHC, endogenous peroxidases were quenched in 0.03% H_2_O_2_ with sodium azide for 5 min. Slides were then incubated in primary antibody (**Table S1**) O/N at RT. Detection was performed by 30 min. incubation with Dako Envision+ System HRP-Labeled Polymers (#K4003, Agilent) followed by a 5-min incubation with DAB. For immunofluorescent staining, slides were blocked in 5% normal goat serum (#01-620-1, ThermoFisher), incubated in primary antibody (**Table S1**) O/N at 4 °C, washed in TBS-T, and incubated in the appropriate secondary antibodies (#A-11011, #A-11077, #A-21131, Invitrogen) for 2 h at RT. EdU was visualized using the Click-iT EdU Cell Proliferation Kit for Imaging (#C10037, ThermoFisher). Nuclear counterstaining was performed using ProLong Gold Antifade Mountant with DAPI (#P36931, ThermoFisher).

### Slide imaging and quantification

Slides were imaged using a Nikon Eclipse E800 microscope with NIS-Elements Basic Research Software or by the Vanderbilt University Medical Center Digital Histology Shared Resource (DHSR). For chromogenic IHC, whole slides were imaged by the DHSR using a Leica SCN400 Slide Scanner (Leica Biosystems). For fluorescent staining, whole slides were imaged by the DHSR using an Aperio Versa 200 automated slide scanner (Leica Biosystems) and visualized using Aperio ImageScope (v12.4.3) (Leica Biosystems). Quantification was performed by an observer blinded to genotype by counting positive cells in well-oriented crypts or crypt-villus units.

### Colon crypt isolation

Murine colon was harvested and splayed longitudinally. The proximal (5 cm from cecum) and distal (5 cm from anus) colon were washed in PBS without calcium or magnesium and minced. Tissue fragments were incubated in chelation buffer (2 mM EDTA in PBS without calcium or magnesium) with nutation for 90 min at 4 °C and washed with PBS followed by 2-min cycles of gentle shaking in shaking buffer (43.3 mM sucrose and 54.9 mM sorbitol in PBS without calcium or magnesium) to free colon crypts. Intact crypts were collected by centrifugation at 150 *x g*, 4 °C for 12 min for RNA isolation and bulk RNA-seq as described below.

### DSS-induced colitis

Bedding was mixed between cages to normalize microbiome 2 wk prior to experiments. Water deprivation caps were placed to accustom mice to drinking from water bottles 1 wk prior to experiments. Water bottles were filled with 2% (w/v) DSS (#DS1004, Gojira Fine Chemicals) for 5 d and then replaced with water for 4 d of recovery prior to sacrifice. At sacrifice, colons were dissected, Swiss-rolled, fixed in 10% NBF, and processed by the Vanderbilt TPSR. H&Es were scored for histologic injury and regeneration by a gastrointestinal pathologist blinded to genotype (MKW) (scoring system described in **Table S2**). Distal colon crypts were collected as described above for RNA isolation and RNA-seq as described below. For EdU experiments, mice were intraperitoneally injected with 50 mg/kg EdU (#NE08701, Biosynth Carbosynth) in PBS 1 h prior to sacrifice. Mice were weighed daily throughout all experiments.

### Inflammatory carcinogenesis

Mice were prepared as described above and injected *i.p.* with 10 mg/kg AOM followed by three 4-day cycles of 3% (w/v) DSS (#DB001, TdB Labs), each followed by 16 d of recovery. Mice were weighed daily. At sacrifice, colons were dissected, flushed with PBS, splayed longitudinally, and imaged using a Nikon SMZ1270 dissecting scope. Tumors were identified by vascular, non-dimpled appearance from these images in conjunction with direct visualization and measured macroscopically in 2 perpendicular dimensions using calipers. Tumor volume was calculated using the previously validated formula W^2^*L/2, where width (W) was the shorter caliper measurement, and length (L) was the longer caliper measurement (85). Tumoroids were generated from a subset of tumors (see **Supplemental Methods**). Histologic injury and regeneration were assessed by MKW (scoring system in **Table S2**). Grade of dysplasia was evaluated as previously described (86) by a gastrointestinal pathologist blinded to genotype (MBP).

### RNA isolation, bulk RNA-seq, and analysis

Murine colon crypts were collected as described above followed by resuspension in TRIzol Reagent (#15596018, Invitrogen) and homogenization by 5x shearing through a 25G needle. Murine tumors were mechanically homogenized in TRIzol using an electronic homogenizer. Samples were centrifuged for 10 min at 16,200 x *g*, 4 °C to remove insoluble material. Cleared homogenate was chloroform-extracted. RNA was isolated from the aqueous layer using the RNeasy Mini Kit (#74106, Qiagen) with on-column DNase treatment (#79254, Qiagen). Library preparation and paired-end 150 bp RNA-seq using the Illumina NovaSeq6000 were performed by Vanderbilt Technologies for Advanced Genomics. Raw reads in FASTQ format were trimmed with fastp (v0.20.0) with default parameters (87). Quantification was performed using Salmon (v1.3.0) (88) against a decoy transcriptome generated from *Mus musculus* GENCODE (v21) (89). Further analysis was performed in R (v3.6.3) as described previously (90). Briefly, quantification files were imported with tximeta (v1.4.5) (91). Genes with counts ≤ 1 were omitted. Differential expression analysis (DEA) was performed on raw transcript counts using DESeq2 (v1.26.0) (92) and annotated with AnnotationDbi (v1.46.1) (93). GSEA (94) was performed on DESeq2-normalized transcript counts using the GSEA software (v4.1.0) with 1,000 gene permutations, gene set randomization, and gene sets found in the Molecular Signatures Database (MSigDB) (v7.2) (95) or derived from the literature (described in **Table S3**). RNA-seq of human CRC and CAC samples collected from 9 regional hospitals in Finland was performed and analyzed as described previously (96) (**Supplemental Methods**).

### *In silico* analyses of publicly available datasets

Raw and processed ChIP-seq data from Baulies *et al*. (16) (#GSE124186) and Murata *et al.* (2) (#GSE130822) were retrieved from the GEO (81). Data were visualized and E box (5’-CATNNG) sequences were annotated using the Broad Institute Integrated Genomics Viewer (v2.9.1) (97). Metadata and SRR files from the PROTECT study (58, 59) (#GSE109142) were downloaded using the NCBI SRA Toolkit and processed as described above for murine samples, with reads quantified against a decoy-aware transcriptome generated from the GENCODE human reference genome (v29) (89). DEA was performed on raw counts using DESeq2 (v1.26.0) (92). Pairwise DEA was also performed on raw counts from RNA-seq of colon biopsies from adults with UC (60) (#GSE128682) and raw counts generated from the TCGA (65) by Rahman *et al*. (64) (#GSE62944) using DESeq2 (v1.33.1) (92). Microarray data from IBD patients (61) (#GSE9686) were analyzed using limma in Geo2R (82). The *Tabula Muris* (41) was queried for *Mtg16* expression using their web interface (https://tabula-muris.ds.czbiohub.org/).

### Statistics

Statistical analyses of human CAC and sporadic colorectal cancer (CRC) samples were performed using limma (v3.34.9) (98). Statistical analyses of all other bulk RNA-seq were performed using DESeq2 (92). Statistical analyses of microarray data were performed using limma in Geo2R (82). *p*_adj_ < 0.05 was considered significant. Statistical analyses for GSEA were performed using the GSEA program (v4.1.0). FDR (q-value) < 0.05 was considered significant. All other statistical analyses were performed in GraphPad Prism (v9.0.1) as indicated in each figure legend. Data are displayed as arithmetic mean ± SEM unless otherwise noted. *p* < 0.05 was considered significant.

### Study approval

All animal experiments were carried out in accordance with protocols approved by the Vanderbilt IACUC. Human CRC and CAC studies (66) were approved by the Ethics Committee of the Hospital District of Helsinki and the Finnish Institute for Health and Welfare.

## Author Contributions

REB designed experiments, performed experiments, and analyzed data. SAA, KMK, CTJ, JMP, and AHG performed experiments and analyzed data. JJ and REB designed and performed bulk RNA-seq data analysis, respectively. KSL, BC, and PNV designed, collected, and analyzed human and murine colon single cell RNA-seq. FR validated antibodies for and performed immunohistochemical staining. MBP and MKW performed histological analyses and provided expertise in gastrointestinal pathology. JJ, RB, and KP analyzed RNA-seq from human tumors. CSW, SPS, SWH, KSL, JAG, KRS, and YAC provided expertise regarding experimental models and experimental design. REB wrote the manuscript. All authors edited and approved the manuscript.

## Acknowledgments

This work was supported by the National Institutes of Health (F30DK120149 to REB, F31DK127687 to PNV, F31CA232272 to JMP, R01DK103831 to KSL, U01CA215798 to KSL, R03DK123489 to JAG, R01CA178030 to SWH, K01DK123495 to SPS, F32DK108492 to SPS, R01DK099204 to CSW, P30DK058404 to the Vanderbilt Digestive Disease Research Center [DDRC], P50CA236733 to the Vanderbilt-Ingram Cancer Center [VICC] Spore in Gastrointestinal Cancer, P30CA068485 and U24DK059637 to the Vanderbilt Translational Pathology Shared Resource [TPSR], and T32GM00734 to the Vanderbilt Medical Scientist Training Program), the U.S. Department of Veterans Affairs Office of Medical Research (1I01BX001426 to CSW and IK2BX004648 to YAC), and the Crohn’s and Colitis Foundation (623541 to CSW and 662877 to SPS). JJ was supported by the Royal Netherlands Academy of Arts and Sciences (Academy Ter Meulen Grant) and the Prince Bernhard Cultural Foundation (Cultural Foundation Grant). KP was supported by grants from Academy of Finland (Finnish Center of Excellence Program 2018–2025, 312041), the iCAN Digital Precision Cancer Medicine Flagship (320185), and the Sigrid Jusélius Foundation.

We thank Barbara Fingleton, Stephen Brandt, Matthew Stier, and other members of the Williams and Hiebert Labs for their critical insights into this manuscript. We would also like to thank the TPSR, Digital Histology Shared Resource (DHSR), Vanderbilt Technologies for Advanced Genomics (VANTAGE), and Advanced Computing Center for Research and Education (ACCRE) for aid with histology, imaging, RNA-seq, and data storage/processing, respectively. Additional computational resources were provided by the ELIXIR node, hosted at the CSC–IT Center for Science, Finland.

## References

1. Beumer J, Clevers H. Cell fate specification and differentiation in the adult mammalian intestine. Nat Rev Mol Cell Bio 2021;22(1):39–53.

2. Murata K et al. Ascl2-Dependent Cell Dedifferentiation Drives Regeneration of Ablated Intestinal Stem Cells. Cell Stem Cell 2020;26(3):377–390.e6.

3. Jadhav U et al. Dynamic Reorganization of Chromatin Accessibility Signatures during Dedifferentiation of Secretory Precursors into Lgr5+ Intestinal Stem Cells. Cell Stem Cell 2017;21(1):65–77.e5.

4. Liu Y, Chen Y-G. Intestinal epithelial plasticity and regeneration via cell dedifferentiation. Cell Regen 2020;9(1):14.

5. Worthington JJ, Reimann F, Gribble FM. Enteroendocrine cells-sensory sentinels of the intestinal environment and orchestrators of mucosal immunity. Mucosal Immunol 2018;11(1):3–20.

6. Skibicka KP, Dickson SL. Enteroendocrine hormones—central effects on behavior. Curr Opin Pharmacol 2013;13(6):977–982.

7. Yamane S, Inagaki N. Control of intestinal stem cell fate: A novel approach to treating diabetes. J Diabetes Invest 2016;7(2):166–168.

8. Parikh K et al. Colonic epithelial cell diversity in health and inflammatory bowel disease. Nature 2019;567(7746):49–55.

9. Aliluev A et al. Diet-induced alteration of intestinal stem cell function underlies obesity and prediabetes in mice. Nat Metabolism 2021;3(9):1202–1216.

10. Chen B, et al. Human colorectal pre-cancer atlas identifies distinct molecular programs underlying two major subclasses of pre-malignant tumors. Biorxiv 2021;2021.01.11.426044.

11. Gelmetti V et al. Aberrant recruitment of the nuclear receptor corepressor-histone deacetylase complex by the acute myeloid leukemia fusion partner ETO. Mol Cell Biol 1998;18(12):7185--91.

12. Cai Y et al. Eto2/MTG16 and MTGR1 are heteromeric corepressors of the TAL1/SCL transcription factor in murine erythroid progenitors. Biochem Bioph Res Co 2009;390(2):295--301.

13. Wang J, Hoshino T, Redner RL, Kajigaya S, Liu JM. ETO, fusion partner in t(8;21) acute myeloid leukemia, represses transcription by interaction with the human N-CoR/mSin3/HDAC1 complex. Proc National Acad Sci 1998;95(18):10860--10865.

14. Steinauer N, Guo C, Zhang J. Emerging Roles of MTG16 in Cell-Fate Control of Hematopoietic Stem Cells and Cancer. Stem Cells Int 2017;2017:1--12.

15. Moore AC et al. Myeloid Translocation Gene Family Members Associate with T-Cell Factors (TCFs) and Influence TCF-Dependent Transcription. Mol Cell Biol 2007;28(3):977–987.

16. Baulies A et al. The Transcription co-Repressors MTG8 and MTG16 Regulate Exit of Intestinal Stem Cells From Their Niche and Differentiation into Enterocyte vs Secretory Lineages. Gastroenterology 2020;159(4):1328–1341.e3.

17. Williams CS et al. MTG16 contributes to colonic epithelial integrity in experimental colitis.. Gut 2013;62(10):1446--55.

18. McDonough EM et al. MTG16 is a tumor suppressor in colitis-associated carcinoma.. Jci Insight 2017;2(16). doi:10.1172/jci.insight.78210

19. Hunt A, Fischer M, Engel ME, Hiebert SW. Mtg16/Eto2 contributes to murine T-cell development. Mol Cell Biol 2011;31(13):2544--51.

20. Acharya P, et al. Mtg16-dependent repression of E protein activity is required for early lymphopoiesis. Biorxiv 2020;2020.07.24.220525.

21. Ghosh HS et al. ETO family protein Mtg16 regulates the balance of dendritic cell subsets by repressing Id2.. J Exp Medicine 2014;211(8):1623--35.

22. Ghosh HS, Liu K, Hiebert S, Reizis B. Eto2/MTG16 Regulates E-Protein Activity and Subset Specification in Dendritic Cell Development. Blood 2012;120(21):1229–1229.

23. Wang L-H, Baker NE. E Proteins and ID Proteins: Helix-Loop-Helix Partners in Development and Disease. Dev Cell 2015;35(3):269--280.

24. Slattery C, Ryan MP, McMorrow T. E2A proteins: Regulators of cell phenotype in normal physiology and disease. Int J Biochem Cell Biology 2008;40(8):1431–1436.

25. Peng V et al. E proteins orchestrate dynamic transcriptional cascades implicated in the suppression of the differentiation of group 2 innate lymphoid cells. J Biol Chem 2020;295(44):14866–14877.

26. Zhuang Y, Cheng P, Weintraub H. B-lymphocyte development is regulated by the combined dosage of three basic helix-loop-helix genes, E2A, E2-2, and HEB.. Mol Cell Biol 1996;16(6):2898–2905.

27. Cisse B et al. Transcription Factor E2-2 Is an Essential and Specific Regulator of Plasmacytoid Dendritic Cell Development. Cell 2008;135(1):37–48.

28. Beck K, Peak MM, Ota T, Nemazee D, Murre C. Distinct roles for E12 and E47 in B cell specification and the sequential rearrangement of immunoglobulin light chain loci. J Exp Med 2009;206(10):2271–2284.

29. Bhattacharya A, Baker NE. A Network of Broadly Expressed HLH Genes Regulates Tissue-Specific Cell Fates. Cell 2011;147(4):881–892.

30. Dréau GL et al. E proteins sharpen neurogenesis by modulating proneural bHLH transcription factors’ activity in an E-box-dependent manner. Elife 2018;7:e37267.

31. Bouderlique T et al. The Concerted Action of E2-2 and HEB Is Critical for Early Lymphoid Specification. Front Immunol 2019;10:455.

32. Rao C, Malaguti M, Mason JO, Lowell S. The transcription factor E2A drives neural differentiation in pluripotent cells. Development 2020;147(12):dev.184093.

33. Banerjee A et al. Succinate Produced by Intestinal Microbes Promotes Specification of Tuft Cells to Suppress Ileal Inflammation. Gastroenterology 2020;159(6):2101–2115.e5.

34. Herring CA et al. Unsupervised Trajectory Analysis of Single-Cell RNA-Seq and Imaging Data Reveals Alternative Tuft Cell Origins in the Gut. Cell Syst 2018;6(1):37–51.e9.

35. Nyström EEL et al. An intercrypt subpopulation of goblet cells is essential for colonic mucus barrier function. Science 2021;372(6539):eabb1590.

36. Poindexter SV et al. Transcriptional corepressor MTG16 regulates small intestinal crypt proliferation and crypt regeneration after radiation-induced injury. Am J Physiol-gastr L 2015;308(6):G562--71.

37. Gunawardene AR, Corfe BM, Staton CA. Classification and functions of enteroendocrine cells of the lower gastrointestinal tract. Int J Exp Pathol 2011;92(4):219–231.

38. Fothergill LJ, Furness JB. Diversity of enteroendocrine cells investigated at cellular and subcellular levels: the need for a new classification scheme. Histochem Cell Biol 2018;150(6):693–702.

39. Akiyama S et al. CCN3 Expression Marks a Sulfomucin-nonproducing Unique Subset of Colonic Goblet Cells in Mice. Acta Histochem Cytoc 2017;50(6):17027.

40. Billing LJ et al. Single cell transcriptomic profiling of large intestinal enteroendocrine cells in mice – Identification of selective stimuli for insulin-like peptide-5 and glucagon-like peptide-1 co-expressing cells. Mol Metab 2019;29:158–169.

41. Schaum N et al. Single-cell transcriptomics of 20 mouse organs creates a Tabula Muris. Nature 2018;562(7727):367–372.

42. Jenny M et al. Neurogenin3 is differentially required for endocrine cell fate specification in the intestinal and gastric epithelium. Embo J 2002;21(23):6338–6347.

43. López-Díaz L et al. Intestinal Neurogenin 3 directs differentiation of a bipotential secretory progenitor to endocrine cell rather than goblet cell fate. Dev Biol 2007;309(2):298–305.

44. Gehart H et al. Identification of Enteroendocrine Regulators by Real-Time Single-Cell Differentiation Mapping. Cell 2019;176(5):1158–1173.e16.

45. Li HJ, Ray SK, Kucukural A, Gradwohl G, Leiter AB. Reduced Neurog3 Gene Dosage Shifts Enteroendocrine Progenitor Towards Goblet Cell Lineage in the Mouse Intestine. Cell Mol Gastroenterology Hepatology [published online ahead of print: 2020]; doi:10.1016/j.jcmgh.2020.08.006

46. Yan KS et al. Intestinal Enteroendocrine Lineage Cells Possess Homeostatic and Injury-Inducible Stem Cell Activity. Cell Stem Cell 2017;21(1):78–90.e6.

47. Engel ME, Nguyen HN, Mariotti J, Hunt A, Hiebert SW. Myeloid translocation gene 16 (MTG16) interacts with Notch transcription complex components to integrate Notch signaling in hematopoietic cell fate specification.. Mol Cell Biol 2010;30(7):1852--63.

48. Flier LG van der, et al. Transcription Factor Achaete Scute-Like 2 Controls Intestinal Stem Cell Fate. Cell 2009;136(5):903–912.

49. Wice BM, Gordon JI. Forced Expression of Id-1 in the Adult Mouse Small Intestinal Epithelium Is Associated with Development of Adenomas*. J Biol Chem 1998;273(39):25310–25319.

50. O’Brien CA et al. ID1 and ID3 Regulate the Self-Renewal Capacity of Human Colon Cancer-Initiating Cells through p21. Cancer Cell 2012;21(6):777–792.

51. Biyajima K et al. Id2 deletion attenuates Apc-deficient ileal tumor formation. Biol Open 2015;4(8):993–1001.

52. Rockman SP et al. Id2 Is a Target of the β-Catenin/T Cell Factor Pathway in Colon Carcinoma*. J Biol Chem 2001;276(48):45113–45119.

53. Yu Y, Yang W, Li Y, Cong Y. Enteroendocrine Cells: Sensing Gut Microbiota and Regulating Inflammatory Bowel Diseases. Inflamm Bowel Dis 2019;26(1):11–20.

54. Khaloian S et al. Mitochondrial impairment drives intestinal stem cell transition into dysfunctional Paneth cells predicting Crohn’s disease recurrence. Gut 2020;69(11):1939–1951.

55. Gersemann M et al. Differences in goblet cell differentiation between Crohn’s disease and ulcerative colitis. Differentiation 2009;77(1):84–94.

56. Sünderhauf A et al. Loss of mucosal p32/gC1qR/HABP1 triggers energy deficiency and impairs goblet cell differentiation in ulcerative colitis. Cell Mol Gastroenterology Hepatology [published online ahead of print: 2021]; doi:10.1016/j.jcmgh.2021.01.017

57. Treveil A, et al. Regulatory network analysis of Paneth cell and goblet cell enriched gut organoids using transcriptomics approaches. Mol Omics 2019;16(1):39–58.

58. Hyams JS et al. Clinical and biological predictors of response to standardised paediatric colitis therapy (PROTECT): a multicentre inception cohort study. Lancet 2019;393(10182):1708–1720.

59. Haberman Y et al. Ulcerative colitis mucosal transcriptomes reveal mitochondriopathy and personalized mechanisms underlying disease severity and treatment response. Nat Commun 2019;10(1):38.

60. Fenton CG, Taman H, Florholmen J, Sørbye SW, Paulssen RH. Transcriptional Signatures That Define Ulcerative Colitis in Remission. Inflamm Bowel Dis 2020;27(1):94–105.

61. Carey R et al. Activation of an IL-6:STAT3-dependent transcriptome in pediatric-onset inflammatory bowel disease. Inflamm Bowel Dis 2008;14(4):446–457.

62. Perse M, Cerar A. Dextran Sodium Sulphate Colitis Mouse Model: Traps and Tricks. J Biomed Biotechnol 2012;2012:1--13.

63. Dieleman, et al. Chronic experimental colitis induced by dextran sulphate sodium (DSS) is characterized by Th1 and Th2 cytokines. Clin Exp Immunol 1998;114(3):385–391.

64. Rahman M et al. Alternative preprocessing of RNA-Sequencing data in The Cancer Genome Atlas leads to improved analysis results. Bioinformatics 2015;31(22):3666–3672.

65. Chang K et al. The Cancer Genome Atlas Pan-Cancer analysis project. Nat Genet 2013;45(10):1113–1120.

66. Rajamäki K et al. Genetic and epigenetic characteristics of inflammatory bowel disease associated colorectal cancer. Gastroenterology [published online ahead of print: 2021]; doi:10.1053/j.gastro.2021.04.042

67. Andrews N et al. An unsupervised method for physical cell interaction profiling of complex tissues. Nat Methods 2021;18(8):912–920.

68. Schonhoff SE, Giel-Moloney M, Leiter AB. Neurogenin 3-expressing progenitor cells in the gastrointestinal tract differentiate into both endocrine and non-endocrine cell types. Dev Biol 2004;270(2):443–454.

69. Wang Y et al. Single-cell transcriptome analysis reveals differential nutrient absorption functions in human intestine. J Exp Med 2019;217(2):e20191130.

70. Beumer J et al. High-Resolution mRNA and Secretome Atlas of Human Enteroendocrine Cells. Cell 2020;181(6):1291–1306.e19.

71. Li HJ, Ray SK, Singh NK, Johnston B, Leiter AB. Basic helix-loop-helix transcription factors and enteroendocrine cell differentiation. Diabetes Obes Metabolism 2011;13(s1):5–12.

72. Longo A, Guanga GP, Rose RB. Crystal Structure of E47−NeuroD1/Beta2 bHLH Domain−DNA Complex: Heterodimer Selectivity and DNA Recognition † , ‡. Biochemistry-us 2008;47(1):218–229.

73. Yui S et al. YAP/TAZ-Dependent Reprogramming of Colonic Epithelium Links ECM Remodeling to Tissue Regeneration. Cell Stem Cell 2018;22(1):35–49.e7.

74. Qu M et al. Establishment of intestinal organoid cultures modeling injury-associated epithelial regeneration. Cell Res 2021;1–13.

75. Lee C-C et al. TCF12 Protein Functions as Transcriptional Repressor of E-cadherin, and Its Overexpression Is Correlated with Metastasis of Colorectal Cancer*. J Biol Chem 2012;287(4):2798–2809.

76. Zhao H et al. E2A suppresses invasion and migration by targeting YAP in colorectal cancer cells. J Transl Med 2013;11(1):317.

77. Short SP et al. Kaiso is required for MTG16-dependent effects on colitis-associated carcinoma. Oncogene 2019;38(25):5091–5106.

78. Chyla BJ et al. Deletion of Mtg16, a Target of t(16;21), Alters Hematopoietic Progenitor Cell Proliferation and Lineage Allocation▿ †. Mol Cell Biol 2008;28(20):6234–6247.

79. Rozenblatt-Rosen O et al. The Human Tumor Atlas Network: Charting Tumor Transitions across Space and Time at Single-Cell Resolution. Cell 2020;181(2):236–249.

80. Liu Q et al. Quantitative assessment of cell population diversity in single-cell landscapes. Plos Biol 2018;16(10):e2006687.

81. Edgar R, Domrachev M, Lash AE. Gene Expression Omnibus: NCBI gene expression and hybridization array data repository. Nucleic Acids Res 2002;30(1):207–210.

82. Barrett T et al. NCBI GEO: archive for functional genomics data sets—update. Nucleic Acids Res 2013;41(D1):D991–D995.

83. Wang F et al. Technical Advance RNAscope A Novel in Situ RNA Analysis Platform for Formalin-Fixed, Paraffin-Embedded Tissues. J Mol Diagnostics 2012;14:22--29.

84. Short SP et al. p120-Catenin is an obligate haploinsufficient tumor suppressor in intestinal neoplasia. J Clin Invest 2017;127(12):4462–4476.

85. Faustino-Rocha A et al. Estimation of rat mammary tumor volume using caliper and ultrasonography measurements. Lab Animal 2013;42(6):217–224.

86. Boivin GP et al. Pathology of mouse models of intestinal cancer: Consensus report and recommendations. Gastroenterology 2003;124(3):762–777.

87. Chen S, Zhou Y, Chen Y, Gu J. fastp: an ultra-fast all-in-one FASTQ preprocessor. Bioinformatics 2018;34(17):i884–i890.

88. Patro R, Duggal G, Love MI, Irizarry RA, Kingsford C. Salmon: fast and bias-aware quantification of transcript expression using dual-phase inference. Nat Methods 2017;14(4):417–419.

89. Frankish A et al. GENCODE reference annotation for the human and mouse genomes. Nucleic Acids Res 2018;47(D1):gky955-.

90. RNA-seq workflow: gene-level exploratory analysis and differential expression [Internet]https://www.bioconductor.org/packages/devel/workflows/vignettes/rnaseqGene/inst/doc/rnaseqGene.html. cited February 5, 2021

91. Love MI et al. Tximeta: Reference sequence checksums for provenance identification in RNA-seq. Plos Comput Biol 2020;16(2):e1007664.

92. Love MI, Huber W, Anders S. Moderated estimation of fold change and dispersion for RNA-seq data with DESeq2. Genome Biol 2014;15(12):550.

93. H P, M C, S F, N L. Bioconductor - AnnotationDbi [Internet]. AnnotationDbi: Manipulation of SQLite-based annotations in Bioconductor. R package version 1.54.1. 2021;https://bioconductor.org/packages/release/bioc/html/AnnotationDbi. cited

94. Subramanian A et al. Gene set enrichment analysis: A knowledge-based approach for interpreting genome-wide expression profiles. P Natl Acad Sci Usa 2005;102(43):15545–15550.

95. Liberzon A et al. Molecular signatures database (MSigDB) 3.0. Bioinformatics 2011;27(12):1739–1740.

96. Rajamäki K et al. Genetic and epigenetic characteristics of inflammatory bowel disease associated colorectal cancer. Gastroenterology [published online ahead of print: 2021]; doi:10.1053/j.gastro.2021.04.042

97. Thorvaldsdóttir H, Robinson JT, Mesirov JP. Integrative Genomics Viewer (IGV): high-performance genomics data visualization and exploration. Brief Bioinform 2013;14(2):178–192.

98. Ritchie ME et al. limma powers differential expression analyses for RNA-sequencing and microarray studies. Nucleic Acids Res 2015;43(7):e47–e47.

