## Supplementary Data for "MTG16 (CBFA2T3) regulates colonic epithelial differentiation, colitis, and tumorigenesis by repressing E protein transcription factors"

### Supplemental Methods

**Tissue microarray (TMA) construction.** For baseline WT and *Mtg16*$^{-/-}$ analyses, the TPSR constructed 2 TMAs using the TMA Grand Master automated arrayer (PerkinElmer) from individual FFPE blocks. Briefly, 2-mm cores were punched from FFPE blocks containing Swiss-rolled WT and *Mtg16*$^{-/-}$ SI and colon (3 cores per Swiss roll). Following construction, H&Es were generated from the TMAs and evaluated for quality before using the TMAs for IHC or other staining.

**Bulk RNA-seq of human CRC and CAC samples.** RNA-seq of human samples collected from 9 regional hospitals in Finland was performed and analyzed as previously described (1). Briefly, RNA was TRIzol-extracted from human tumor samples and underwent HiSeq LncRNA-Seq library preparation and paired-end sequencing using Illumina HiSeqXTen. Raw sequences were mapped onto the human transcriptome (Ensembl release 79) using Salmon (v0.12.0) (2). Gene-level quantification with variance stabilization was performed using DESeq2 (v1.18.1) (3) followed by limma (v3.34.9) correction of sequencing batch effects (4).

**AOM/DSS tumoroid cultures.** Colon tumors were isolated at sacrifice following the AOM/DSS protocol. Tumoroids were generated as previously described (5). Briefly, tumors were diced and incubated in 10 mL digestion buffer (0.1 mg/mL collagenase XI [#C7657, Sigma] and 0.125 mg/mL dispase II [#17105-041, Gibco] in complete Advanced DMEM [#12491015, Gibco]) at 37 °C on a nutating shaker for 2 h. Epithelial cells were mechanically dissociated by pipetting. Supernatant containing the epithelial cells was centrifuged at 150 *x g* for 3 min at 4 °C, washed twice in ice-cold PBS, and plated in 50-µL Matrigel (#356231, Corning) plugs overlaid with 500 µL MGM-CM (minigut media with R-spondin and Noggin conditioned media, generated as previously described (5)) containing 0.002% primocin (#NC9141851, Invivogen). Tumoroids were passaged by gentle dissociation in TrypLE (#12604013, Gibco) at 37 °C, washing in ice-cold PBS, and replating at least once to remove debris before tumoroid collection. Tumoroid fixation and embedding in agarose for staining was then performed as previously described (5).

**Data visualization and presentation.** Figures were generated using BioRender (https://biorender.com/), GSEA (v4.1.0), the R packages Seurat (v4.0.4), pheatmap (v1.0.12), RColorBrewer (v1.1-2), and ggplot2 (v3.2.1), Meta-Chart (https://www.meta-chart.com/venn#/display), the Broad Institute Integrated Genomics Viewer (6) (v2.9.1), GraphPad Prism (v9.0.1), the *Tabula Muris* (7) web interface (https://tabula-muris.ds.czbiohub.org/), and Inkscape (v1.0.1).

**Code availability.** All code is available upon reasonable request.

### Supplemental Tables

| | Primary Antibody | Catalog # | Supplier | Species/ Isotype | Dilution | Antigen retrieval |
|---|---|---|---|---|---|---|
| Chromogenic IHC | α-SYP | ab32127 | Abcam | Rabbit monoclonal [YE269] | 1:800 | Tris-EDTA pH 9.0 in a pressure cooker at 97 °C for 15 min followed by 10 min at RT |
| | α-DCLK1 | 62257 | Cell Signaling Technologies (CST) | Rabbit monoclonal (IgG) | 1:500 | 10 mM sodium citrate, pH 6.0 in a pressure cooker at 105 °C for 15 min followed by 10 min at RT |
| Immunofluorescent staining | α-E-cadherin | 610182 | BD Biosciences | Mouse monoclonal [36] ($IgG_{2a}$) | 1:500 | 10 mM sodium citrate, pH 6.0 at 95 °C for 10 min followed by 30 min cooling at RT |
| | α-CHGA | 20085 | ImmunoStar | Rabbit polyclonal | 1:2000 | 10 mM sodium citrate, pH 6.0 at 95 °C for 10 min followed by 30 min cooling at RT |
| | α-p-H3 (Ser 10) | 06-570 | Sigma-Aldrich | Rabbit polyclonal | 1:400 | 10 mM sodium citrate, pH 6.0 at 95 °C for 10 min followed by 30 min cooling at RT |
| | α-CC3 (Asp175) | 9661 | CST | Rabbit polyclonal | 1:400 | 10 mM sodium citrate, pH 6.0 at 95 °C for 10 min followed by 30 min cooling at RT |
| | α-ɣH2A.X (Ser139) | 9718 | CST | Rabbit monoclonal [20E3] (IgG) | 1:400 | 10 mM sodium citrate, pH 6.0 at 95 °C for 10 min followed by 30 min cooling at RT |
| | α-Ly6B.2 | MCA771G | Bio-Rad | Rat monoclonal ($IgG_{2a}$) | 1:200 | 10 mM sodium citrate, pH 6.0 at 95 °C for 10 min followed by 30 min cooling at RT |
| | α-F4/80 | MCA497G | Bio-Rad | Rat monoclonal [CI:A3-1] ($IgG_{2b}$) | 1:400 | 20 µg/mL proteinase K in TE buffer, pH 8.0 for 3 min at RT |
| | α-Ki67 | Ab15580 | Abcam | Rabbit polyclonal | 1:500 | 10 mM sodium citrate, pH 6.0 at 95 °C for 10 min followed by 30 min cooling at RT |

**Table S1. Antibodies and antigen retrieval used in chromogenic and immunofluorescent staining.**

| Category | Description | Score |
|---|---|---|
| Inflammation (0-3) | None | 0 |
| | Slight | 1 |
| | Moderate | 2 |
| | Severe | 3 |
| % involved by inflammation (1-4) | 1-25% | 1 |
| | 26-50% | 2 |
| | 51-75% | 3 |
| | 76-100% | 4 |
| Depth of inflammation (0-3) | None | 0 |
| | Mucosal | 1 |
| | Mucosal and submucosal | 2 |
| | Transmural | 3 |
| Crypt damage (1-4) | Basal 1/3 crypt cells damaged | 1 |
| | Basal 2/3 crypt cells damaged | 2 |
| | Only surface epithelium intact | 3 |
| | Entire crypt and epithelial surface lost | 4 |
| % involved by crypt damage (1-4) | 1-25% | 1 |
| | 26-50% | 2 |
| | 51-75% | 3 |
| | 76-100% | 4 |
| Epithelial regeneration (0-3) | Tissue appears normal (no residual injury/ complete epithelial regeneration) | 0 |
| | Slight epithelial injury with almost complete regeneration | 1 |
| | Surface epithelium not intact (regeneration present, but epithelial integrity has not been restored) | 2 |
| | Ulcer with no regeneration/tissue repair | 3 |
| Crypt distortion and branching (0-3) | Normal crypts | 0 |
| | Mild | 1 |
| | Moderate | 2 |
| | Severe (mucosa unable to regenerate normal crypt architecture) | 3 |

**Table S2. Histologic injury and regeneration scoring system.** Adapted from Dieleman *et al*. (8) and Fukata *et al*. (9). Note that a higher epithelial regeneration score represents a decreased ability to regenerate.

| Figure(s) | Gene set name | Description | Source |
|---|---|---|---|
| Differentiated epithelial cells (**Figs. 2B, 3F**) | Enteroendocrine (EE) | Expression signature defined for enteroendocrine cells following scRNA-seq and unbiased clustering of FACS-sorted murine intestinal epithelium | Haber *et al.* 2017 (10) |
| | L-cell | Genes enriched in L-cell cluster generated from droplet-based scRNA-seq of healthy human colon mucosa | Parikh *et al.* 2019 (11) |
| | Enterochromaffin | Genes enriched in enterochromaffin cluster generated from droplet-based scRNA-seq of healthy human colon mucosa | Parikh *et al.* 2019 (11) |
| | Absorptive | Expression signature defined for absorptive enterocytes following scRNA-seq and unbiased clustering of FACS-sorted murine intestinal epithelium | Haber *et al.* 2017 (10) |
| | Tuft | Expression signature defined for tuft cells following scRNA-seq and unbiased clustering of FACS-sorted murine intestinal epithelium | Haber *et al.* 2017 (10) |
| | Goblet | Expression signature defined for goblet cells following scRNA-seq and unbiased clustering of FACS-sorted murine intestinal epithelium | Haber *et al.* 2017 (10) |
| | BEST4/OTOP2$^+$ colonocyte | Genes enriched in a novel BEST4/OTOP2$^+$ colonocyte cluster generated from droplet-based scRNA-seq of healthy human colon mucosa | Parikh *et al.* 2019 (11) |
| Enteroendocrine (EE) progenitor cells (**Figs. 2B, 3F**) | Early EE | Genes enriched in FACS-sorted early EE progenitors identified using a novel “Neuorg3Chrono” pulse-chase reporter mouse that allows temporal resolution of cells in the EE lineage | Gehart *et al.* 2019 (12) |
| | Early/Intermediate EE | Genes enriched in FACS-sorted early and intermediate EE progenitors identified using a novel “Neuorg3Chrono” pulse-chase reporter mouse that allows temporal resolution of cells in the EE lineage | Gehart *et al.* 2019 (12) |
| | Intermediate EE | Genes enriched in FACS-sorted intermediate EE progenitors identified using a novel “Neuorg3Chrono” pulse-chase reporter mouse that allows temporal resolution of cells in the EE lineage | Gehart *et al.* 2019 (12) |
| | Intermediate/Late EE | Genes enriched in FACS-sorted intermediate and late EE progenitors identified using a novel “Neuorg3Chrono” pulse-chase reporter mouse that allows temporal resolution of cells in the EE lineage | Gehart *et al.* 2019 (12) |
| | Late EE | Genes enriched in FACS-sorted late EE progenitors identified using a novel “Neuorg3Chrono” pulse-chase reporter mouse that allows temporal resolution of cells in the EE lineage | Gehart *et al.* 2019 (12) |
| | *Neurog3*-EGFP$^{+/+}$ | Genes enriched in FACS-sorted *Neurog3*-EGFP$^+$ intestinal epithelial cells from mice homozygous for the *Neurog3*-EGFP reporter | Li *et al.* 2020 (13) |

| | | | |
|---|---|---|---|
| Stem cells and progenitor cells (**Figs. 2C, 3G**) | *Bmi1*-GFP$^{+}$ | Genes enriched in FACS-sorted *Bmi1*-GFP$^{+}$ murine intestinal epithelial cells | Yan *et al.* 2017 (14) |
| | *mTert*-GFP$^{+}$ | Genes enriched in FACS-sorted *mTert*-GFP$^{+}$ murine intestinal epithelial cells | Yan *et al.* 2017 (14) |
| | FVR$^{Low}$ | Genes enriched in FACS-sorted *Fltp*-H2B-Venus reporter (FVR)$^{Low}$ murine intestinal epithelial cells. FVR$^{Low}$ cells were characterized as quiescent, terminally differentiated EE cells that had previously induced *Fltp* expression (activation of the WNT-PCP pathway). | Bottcher *et al.* 2021 (15) |
| | *Rbpj*-DBZ Sec-Pro | Genes enriched in intestinal crypts from both *Rbpj*$^{-/-}$ and dibenzazepine(DBZ)-treated mice (2 independent ways of inhibiting Notch signaling to increase secretory progenitors [Sec-Pro]) | Kim *et al.* 2014 (16) |
| | CD166$^{Hi}$ | Genes enriched in FACS-sorted CD166$^{Hi}$ (antibody-based) murine intestinal epithelial cells | Yan *et al.* 2017 (14) |
| | *MKI67*$^{Hi}$ ("GAO_LARGE_INTESTINE_ADULT_CH_MKI67HIGH_CELLS" in the MSigDB) | Genes enriched in *MKI67*$^{Hi}$ cell cluster from scRNA-seq of human colon from healthy adults | Gao *et al.* 2018 (17); available in the MSigDB (18) |
| | LRIG1$^{+}$ | Genes enriched in FACS-sorted LRIG1$^{+}$ (antibody-based) murine intestinal epithelial cells | Powell *et al.* 2012 (19) |
| | *Lgr5*$^{+}$ ISC | See "Munoz2012_Lgr5-GFPHigh_ISC" (*Lgr5*$^{+}$ stem cell gene sets). | Muñoz *et al*. 2012 (20) |
| | *Lgr5*$^{+}$ ISC (colon) | See "Murata2020_Lgr5_Stem-Cell_Colon_High" (*Lgr5*$^{+}$ stem cell gene sets). | Murata *et al.* 2020 (21) |
| *Lgr5*$^{+}$ stem cell gene sets (**Fig. S6A**) | Yan2017-ISC-Lgr5-Cre_Pos_UP | Genes enriched in FACS-sorted *Lgr5*-eGFP$^{+}$ murine intestinal epithelial cells | Yan *et al.* 2017 (14) |
| | Basak2017_UP-in-active-cycling-Lgr5_Pos | Genes enriched in FACS-sorted *Lgr5*-eGFP$^{+}$ *Ki67*-RFP$^{+}$ murine intestinal epithelial cells | Basak *et al.* 2017 (22); derived from Basak *et al.* 2014 (23) |
| | Basak2017_UP-in-quiescent-Lgr5_Pos | Genes enriched in FACS-sorted *Lgr5*-eGFP$^{+}$ *Ki67*-RFP$^{-}$ murine intestinal epithelial cells | Basak *et al.* 2017 (22); derived from Basak *et al.* 2014 (23) |

| | | | |
|---|---|---|---|
| | Munoz2012_Lgr5-GFPHigh_ISC | ISC signature derived from complementary transcriptomic (using multiple microarray platforms) and proteomic profiling of FACS-sorted *Lgr5*-eGFP$^{Hi}$ cells. This gene set was chosen for the main figures (**Fig. 2C, 3G**) because it is widely used in the literature to represent *Lgr5*$^{+}$ ISCs. | Muñoz *et al*. 2012 (20) |
| | Murata2020_Lgr5_Stem-Cell_Colon_Total | Out of the genes specific to *Lgr5*$^{+}$ ISCs according to Muñoz *et al*.(20), genes expressed at any level (> 1 RPKM) in FACS-sorted *Lgr5*$^{Dtr\text{-}GFP}$ cells from uninjured murine colon (401 genes) | Murata *et al.* 2020 (21) |
| | Murata2020_Lgr5_Stem-Cell_Colon_High | Out of the genes specific to *Lgr5*$^{+}$ ISCs according to Muñoz *et al*.(20), genes expressed at "appreciable levels" (> 10 RPKM) in FACS-sorted *Lgr5*$^{Dtr\text{-}GFP}$ cells from uninjured murine colon (176 genes). This gene set was chosen for the main figures (**Fig. 2C, 3G**) because it was generated recently using colonic *Lgr5*$^{+}$ stem cells. | Murata *et al.* 2020 (21) |
| | Yan2017-SIclusters_C1_non-cycling-Lgr5-GFP | Genes enriched in cluster representing non-cycling *Lgr5*$^{+}$ ISCs following scRNA-seq of FACS-isolated *Lgr5*-eGFP$^{+}$ cells, *Bmi1*-GFP$^{+}$ cells, and *Prox1*-GFP$^{+}$ cells versus a fourth control sample of *Lgr5*-eGFP$^{-}$ intestinal epithelial cells | Yan *et al.* 2017 (14) |
| | GAO_LARGE_INTESTINE_24W_C5_LGR5POS_STEM_CELL | Genes enriched in *Lgr5*$^{+}$ stem cell cluster from scRNA-seq of human fetal large intestine | Gao *et al.* 2018 (17); available in the MSigDB (18) |
| Intestinal epithelial signaling pathways (**Fig. S6B**) | GO_CANONICAL_WNT_SIGNALING_PATHWAY | | MSigDB (18) (Gene Ontology [GO]) |
| | GO_EPIDERMAL_GROWTH_FACTOR_RECEPTOR_SIGNALING_PATHWAY | | MSigDB (18) (GO) |
| | GO_HIPPO_SIGNALING | | MSigDB (18) (GO) |
| | GO_NON_CANONICAL_WNT_SIGNALING_PATHWAY | | MSigDB (18) (GO) |
| | GO_NOTCH_SIGNALING_PATHWAY | | MSigDB (18) (GO) |
| | HALLMARK_KRAS_SIGNALING_UP | | MSigDB (18) (Hallmarks) |
| | KEGG_HEDGEHOG_SIGNALING_PATHWAY | | MSigDB (18) (Kyoto Encyclopedia of Genes and Genomes [KEGG]) |
| | KEGG_MAPK_SIGNALING_PATHWAY | | MSigDB (18) (KEGG) |
| | KEGG_TGF_BETA_SIGNALING_PATHWAY | | MSigDB (18) (KEGG) |

| | | | |
|---|---|---|---|
| | PID_BMP_PATHWAY | | MSigDB (18) (Protein Interactions Database [PID]) |
| | PID_NOTCH_PATHWAY | | MSigDB (18) (PID) |
| | PID_RAS_PATHWAY | | MSigDB (18) (PID) |
| | WNT-PCP_PATHWAY | | Compiled from Bottcher *et al.* 2021 (15) and Smith *et al.* 2020 (24) |
| E and ID protein target genes (**Figs. 2E, 3H, 5J-K**) | E2-2 (“ TCF4_Q5” in the MSigDB) | E2-2 transcription factor target genes (Note: this is different from the WNT effector TCF4) | MSigDB (18) |
| | E47_01 | E47 (splice isoform of E2A) transcription factor target genes | MSigDB (18) |
| | E47_02 | E47 (splice isoform of E2A) transcription factor target genes | MSigDB (18) |
| | E12_Q6 | E12 (splice isoform of E2A) transcription factor target genes | MSigDB (18) |
| | HEB_Q6 | HEB transcription factor target genes | MSigDB (18) |
| | E2A_Q2 | E2A transcription factor target genes | MSigDB (18) |
| | ID1_TARGETS (ID1_TARGET_GENES in the MSigDB) | Genes associated with upregulation of ID1 | Kolmykov *et al.* 2020 (25); available in the MSigDB (18) |
| | ID2_TARGETS (ID2_TARGET_GENES in the MSigDB) | Genes associated with upregulation of ID2 | Kolmykov *et al.* 2020 (25); available in the MSigDB (18) |
| Intestinal epithelial regeneration (**Figs. 5H-I, 6G-H**) | DSS-induced colitis regenerating epithelium | Genes enriched in regenerating colonic epithelium following DSS-induced injury | Wang *et al.* 2019 (26); derived from Yui *et al.* 2018 (27) |
| | Fetal epithelial gene signature | Genes enriched in fetal epithelial spheroids and “fetal-like reprogramming” of the regenerating intestinal epithelium | Wang *et al.* 2019 (26); derived from Mustata *et al.* 2013 (28) and Yui *et al.* 2018 (27) |
| | scRNA-seq cluster SSC2c “revival” cells | Genes enriched in sub-cluster SSC2c generated from unsupervised clustering of scRNA-seq generated from 12 Gy-irradiated and non-irradiated mouse intestinal epithelium. The SSC2c cluster, which appeared only after irradiation, was reported to contain quiescent “revival” stem cells induced by the YAP1 transcription factor. | Ayyaz *et al.* 2019 (29) |

| | | | |
|---|---|---|---|
| | *Clu*$^{+}$ “revival” cell signature | *Clu*-GFP$^{+}$ cells FACS-sorted from irradiated *BAC-Clu*-GFP mice, described as a damage-induced “revival” stem cell population | Ayyaz *et al.* 2019 (29) |
| | Composite injury-regeneration signature | Gene set compiled from the above DSS-induced colitis regenerating epithelial, fetal epithelial, and “revival” cell signatures | Qu *et al.* 2021 (30) |
| | *Ascl2*$^{+}$ de-differentiating cell signature | Genes significantly upregulated in FACS-sorted, de-differentiating/regenerating *Ascl2*-mCh$^{+}$ “upper” cells from murine colon following diphtheria toxin (DT)-mediated ablation of Lgr5$^{+}$ stem cells (compared to *Lgr5*$^{+}$ stem cells FACS-sorted from uninjured $Lgr5^{Dtr\text{-}GFP}$ mouse colon) | Murata *et al.* 2020 (21) |

**Table S3. Gene sets used for GSEA.**

### Supplemental Figures

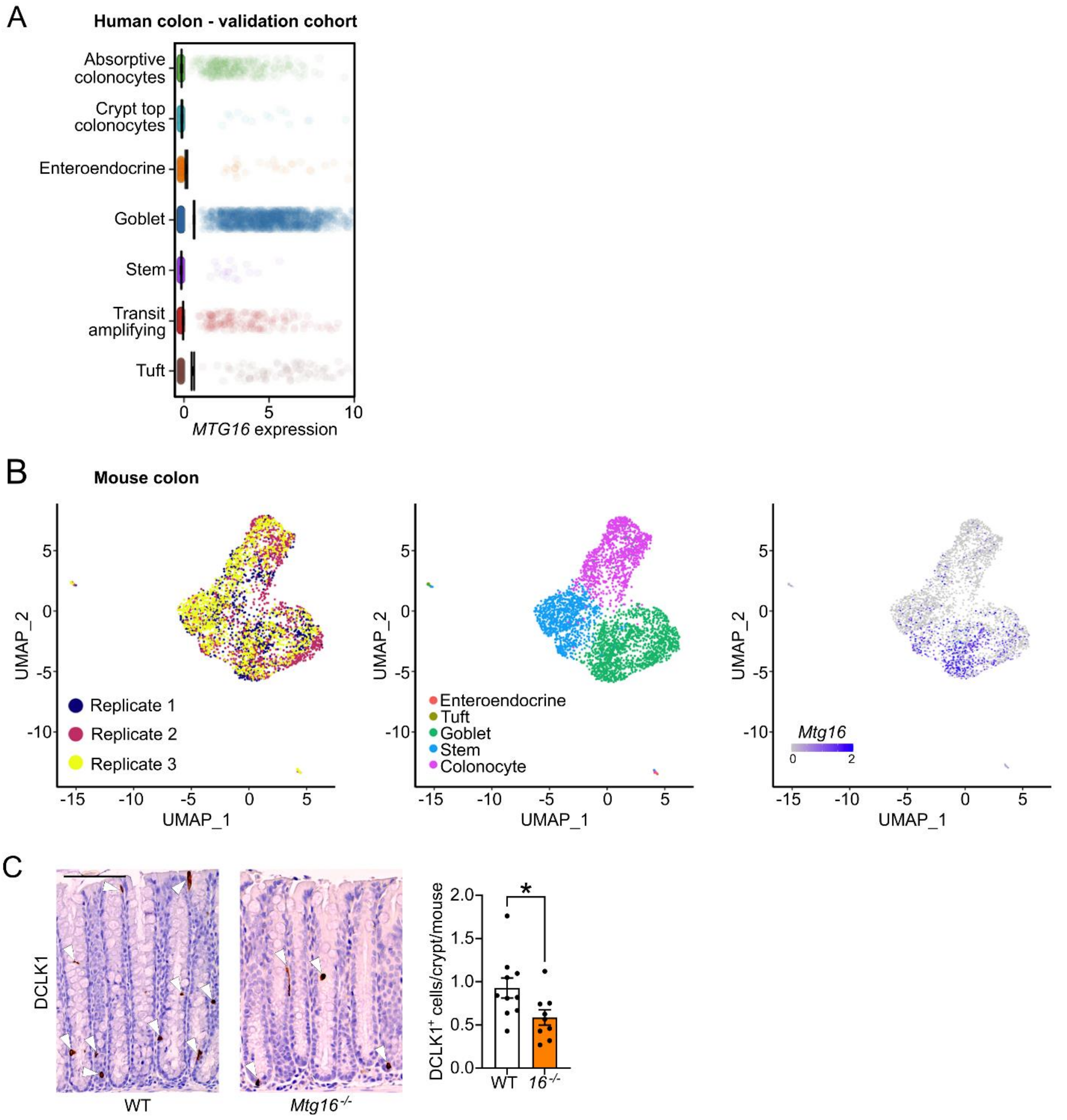


**Figure S1. Data related to Fig. 1. (A)** *MTG16* expression in scRNA-seq of the human colon (validation cohort). 34,008 cells were sequenced from n = 31 normal human colon samples. **(B)** *Mtg16* expression in scRNA-seq of murine colonic epithelial isolates summarized by cell type in Fig. 1B. Left, UMAP plot demonstrating n = 3 biological replicates. Middle, UMAP plot annotated by cell type. Right, UMAP plot demonstrating *Mtg16* expression in individual cells. 3,653 cells were sequenced. **(C)** WT and *Mtg16*$^{-/-}$ mouse colon (n = 10 WT, 9 *Mtg16*$^{-/-}$) stained for tuft cells by IHC for doublecortin-like kinase 1 (DCLK1). Representative images at left. DCLK1$^{+}$ cells are indicated with white arrowheads. Scale bar = 100 µm. *$p < 0.05$ by Mann-Whitney test.

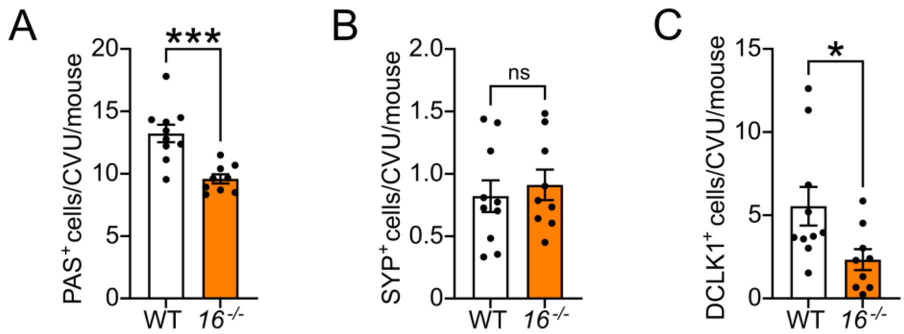


**Figure S2. Secretory cell frequencies in the *Mtg16*$^{-/-}$ SI.** Quantification of **(A)** goblet cells per crypt-villus unit (CVU) by periodic acid-Schiff (PAS) stain, **(B)** enteroendocrine cells by IHC for synaptophysin (SYP), and **(C)** tuft cells by IHC for doublecortin-like kinase 1 (DCLK1) in WT and *Mtg16*$^{-/-}$ small intestine (n = 10 WT, 9 *Mtg16*$^{-/-}$). **p* < 0.05, ****p* < 0.001 by Mann-Whitney test.

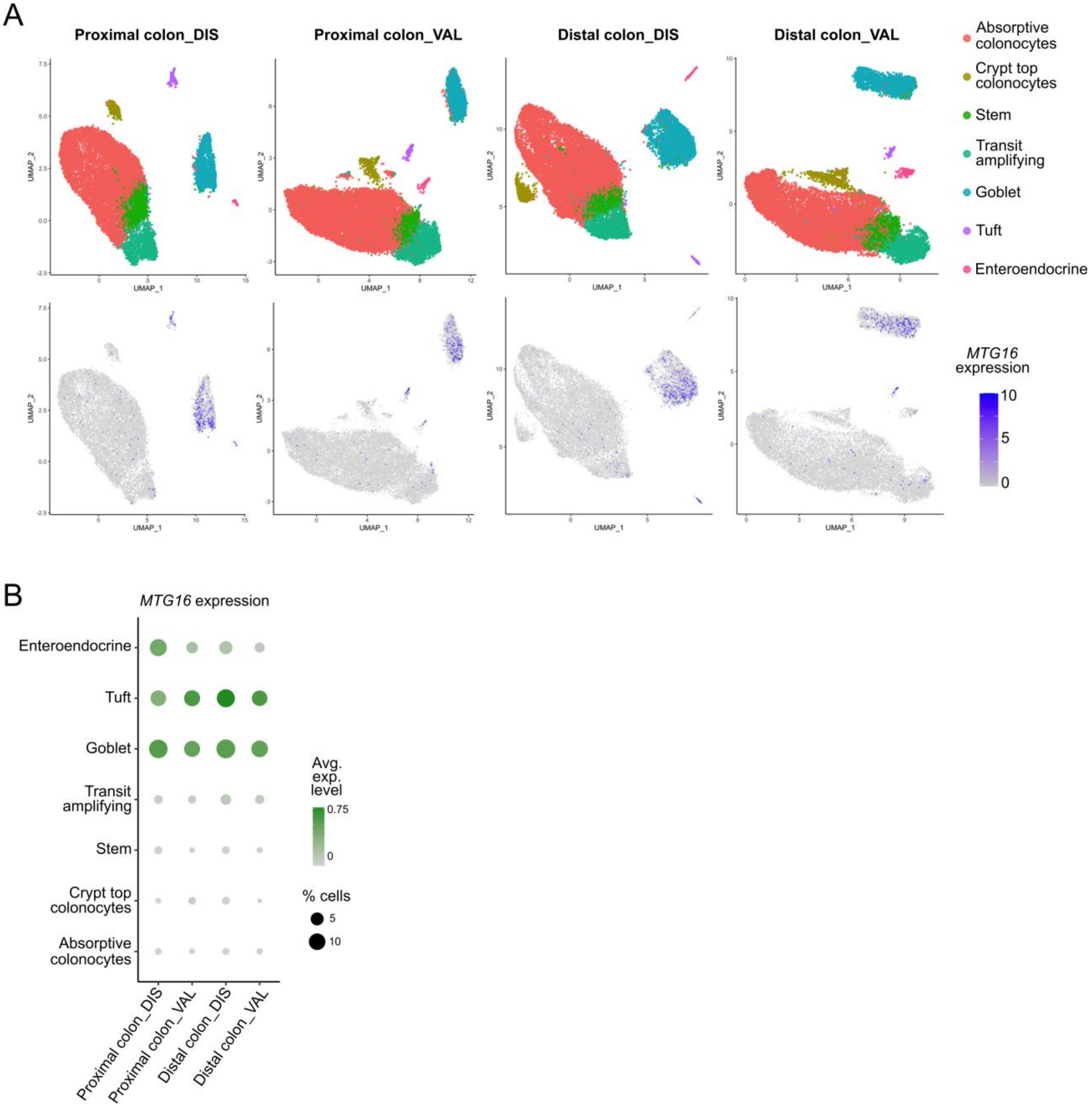


**Figure S3. Differences between the proximal and distal human colon. (A)** Annotated colonic epithelial clusters (top) and corresponding *MTG16* expression (bottom) in human proximal and distal normal colon biopsies queried from our scRNA-seq discovery (DIS) (n = 12,596 cells sequenced from 18 proximal colon samples and 17,778 cells sequenced from 17 distal colon samples) and validation (VAL) (n = 17,289 cells sequenced from 17 proximal colon samples and 16,719 cells sequenced from 14 distal colon samples) cohorts. **(B)** *MTG16* expression in the proximal and distal colon of each human cohort. Color gradient represents the average *MTG16* expression level in each cell population. Dot size represents the percentage of cells in each population expressing *MTG16.*

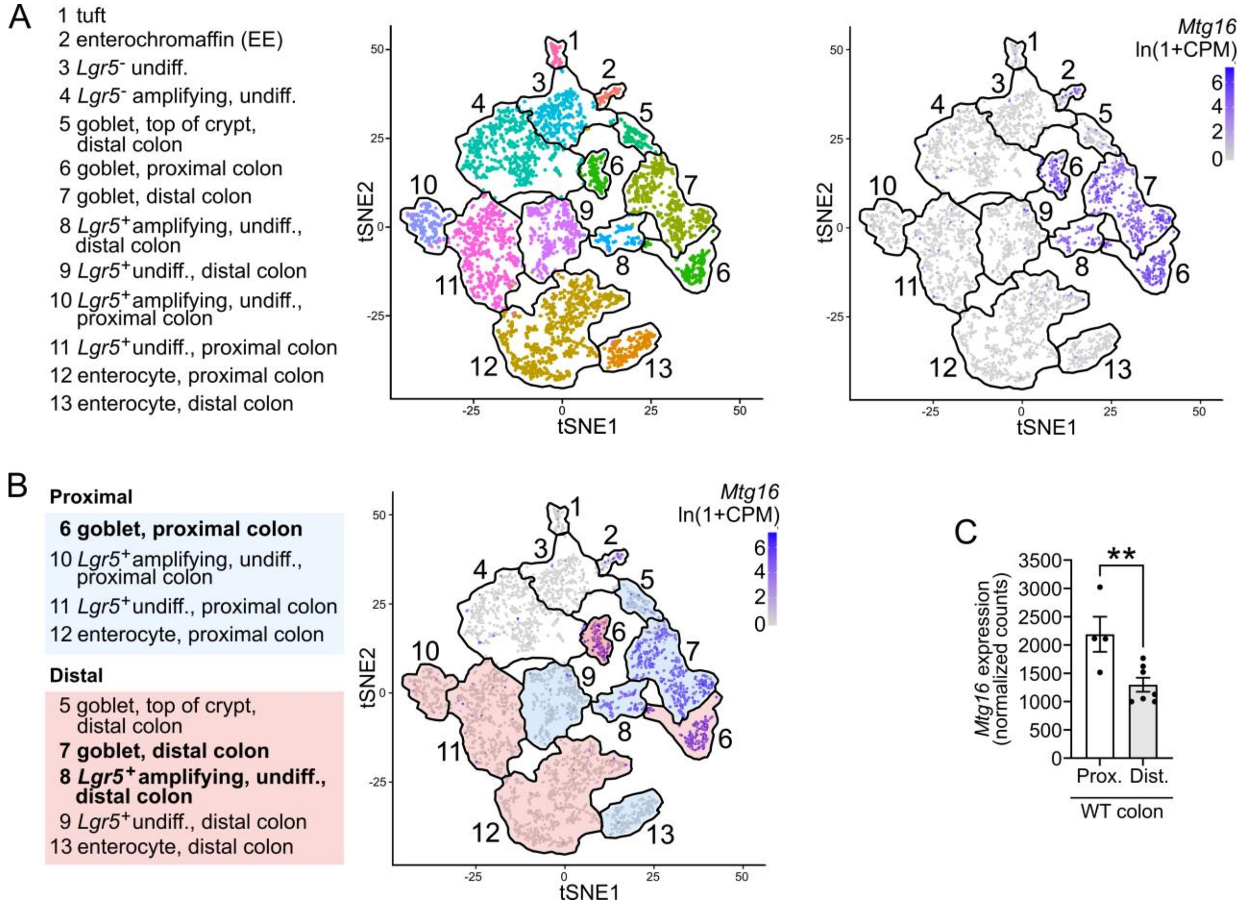


**Figure S4. Differences between the proximal and distal mouse colon. (A)** Annotated colonic epithelial clusters (left) and corresponding *Mtg16* expression (right) queried from scRNA-seq of WT mouse colon (n = 6) publicly available in the *Tabula Muris* (7). **(B)** Alternate annotation emphasizing clusters representing cells specifically from the proximal (blue) or distal (red) colon. Bolded clusters denote clusters expressing *Mtg16*. **(C)** *Mtg16* expression (transcript counts normalized by DESeq2) in WT proximal and distal colon epithelial isolates (n = 4 proximal, 7 distal). **$p_{adj} < 0.01$ by DESeq2.

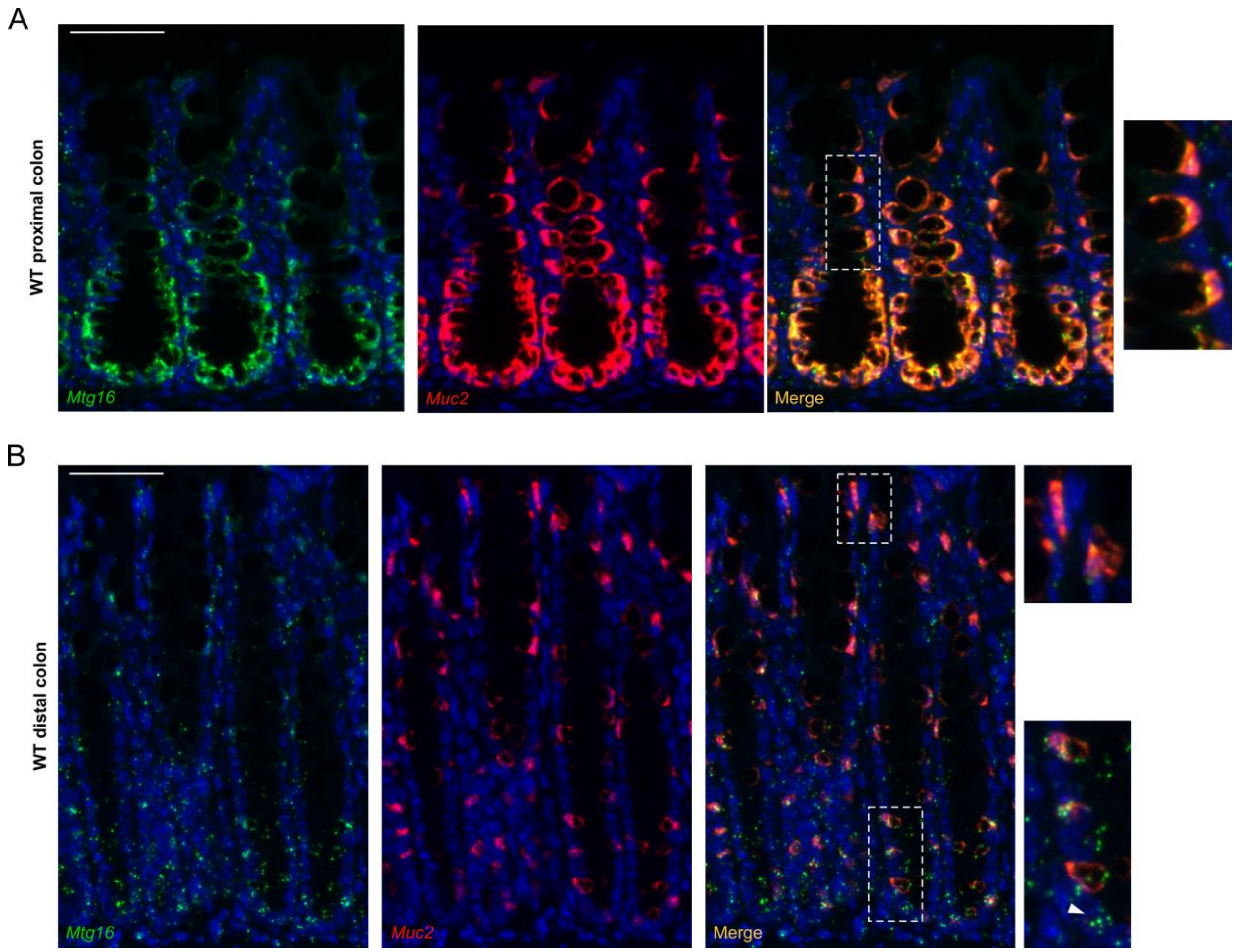


**Figure S5. RNAscope *in situ* hybridization of *Mtg16* and *Muc2* indicating differential expression and co-localization of mRNA expression in WT mouse proximal (A) and distal (B) colon.** Scale bars = 50 µm. White dashed lines denote insets at right. White arrow in **(B)** denotes an epithelial cell near the crypt base expressing *Mtg16*, but not *Muc2*.

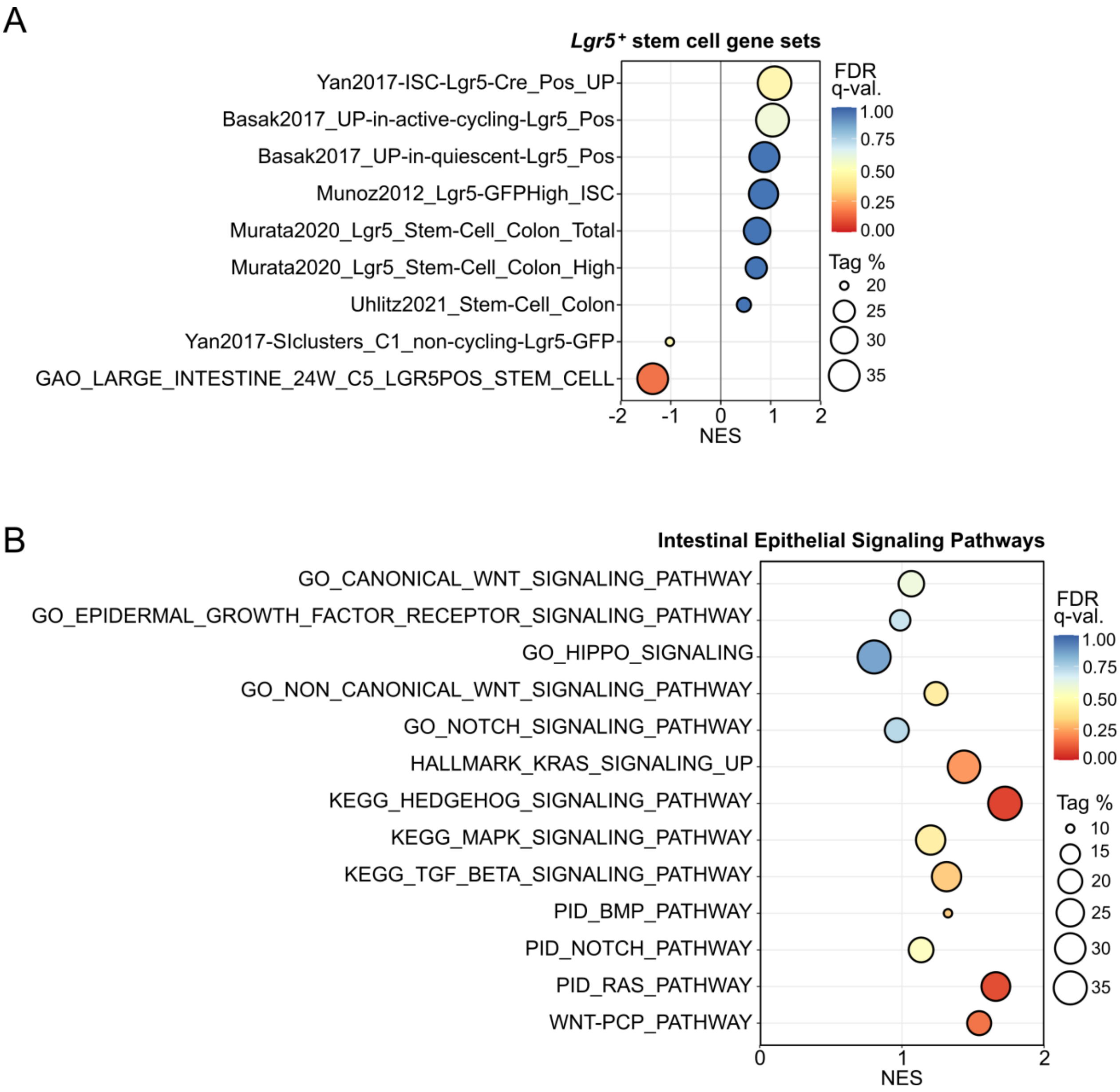


**Figure S6. GSEA of stem cell and signaling pathway gene sets in the *Mtg16$^{-/-}$* distal colon.** GSEA of distal colon RNA-seq (n = 4 WT, 4 *Mtg16$^{-/-}$*) using **(A)** multiple gene sets representing *Lgr5$^{+}$* stem cells derived from the literature and **(B)** gene sets for intestinal epithelial signaling pathways (described in Table S3). NES, normalized enrichment score (ES). Tag % is defined as the percentage of gene hits before (for positive ES) or after (for negative ES) the peak in the running ES, indicating the percentage of genes contributing to the ES. FDR q-value < 0.05 is considered significant.

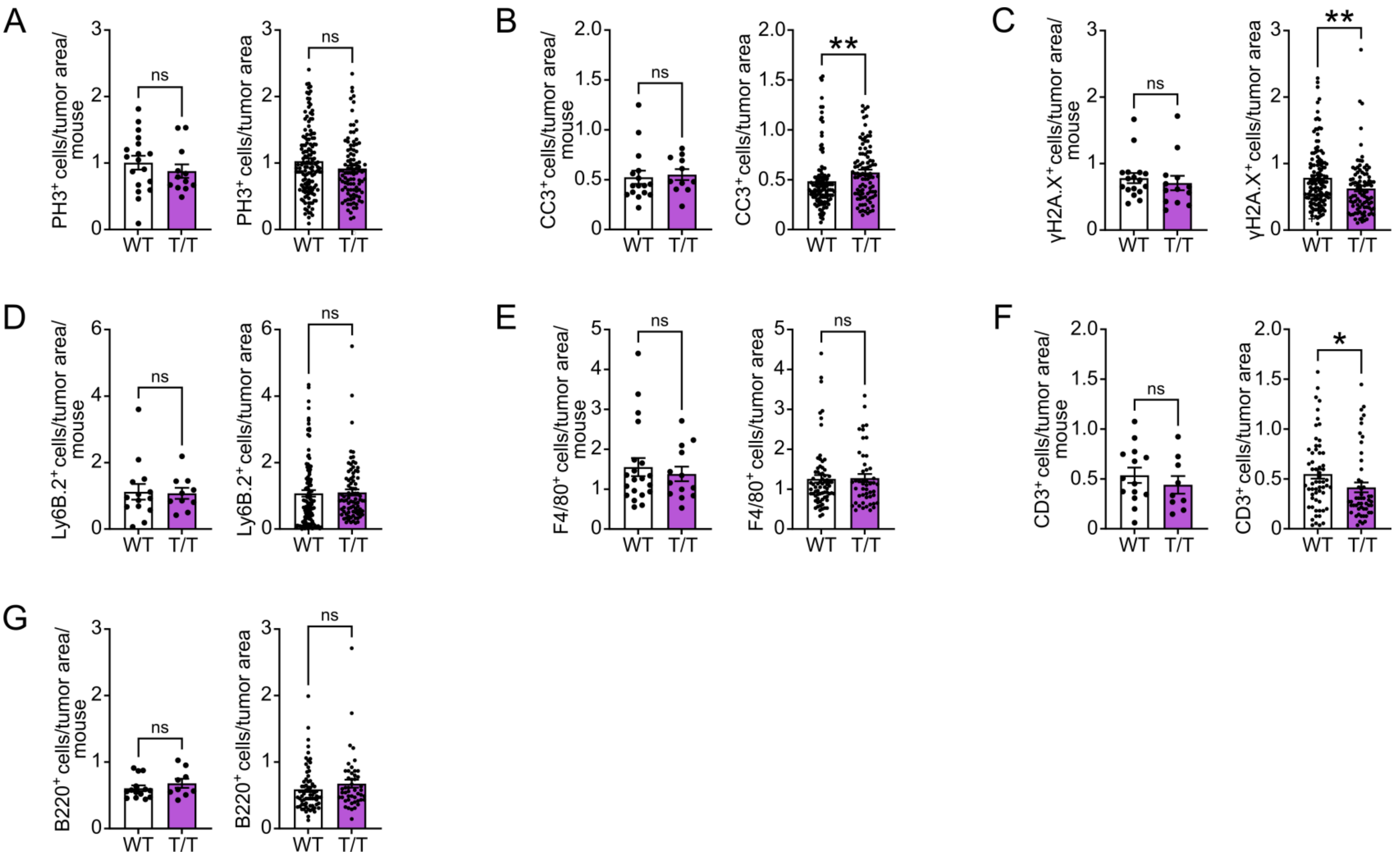


**Figure S7. Additional characterization of AOM/DSS tumors by immunofluorescent staining.** Characterization of tumor epithelial cells by quantification of **(A)** phospho-histone H3-positive (PH3$^+$) proliferating cells, **(B)** cleaved caspase-3-positive (CC3$^+$) cells undergoing apoptosis, and **(C)** γH2A.X$^+$ cells (nuclei) displaying DNA damage. Quantification of intratumoral **(D)** Ly6B.2$^+$ neutrophils, **(E)** F4/80$^+$ macrophages, **(F)** CD3$^+$ T cells, and **(G)** B220$^+$ B cells. **(A-G)** Left, quantification and analysis by average number of cells per tumor area (10^6 pixels) per mouse (n = 9-18 mice). Right, quantification analyzed by tumor area (n = 48-137). *$p$ < 0.05, **$p$ < 0.01 by Mann-Whitney test.

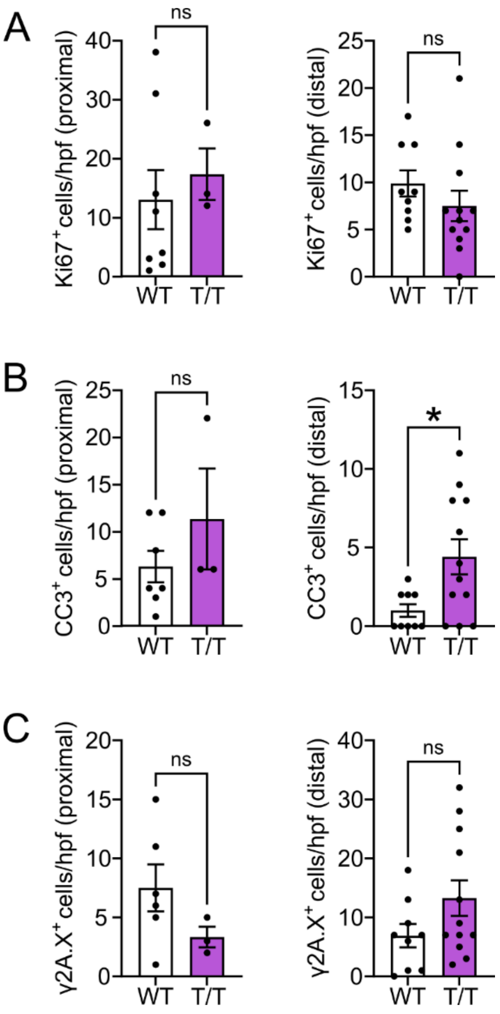


**Figure S8. Characterization of tumoroids derived from proximal (left) and distal (right) AOM/DSS tumors by immunofluorescent staining.** Quantification of **(A)** Ki67$^+$ proliferating cells, **(B)** cleaved caspase-3-positive (CC3$^+$) cells undergoing apoptosis, and **(C)** yH2A.X$^+$ cells (nuclei) displaying DNA damage per high-power field (hpf). n = 3-12 hpf from 1-4 tumoroid lines. *$p$ < 0.05 by Mann-Whitney test.

### Supplemental References


1. Rajamäki K et al. Genetic and epigenetic characteristics of inflammatory bowel disease associated colorectal cancer. *Gastroenterology* [published online ahead of print: 2021]; doi:10.1053/j.gastro.2021.04.042

2. Patro R, Duggal G, Love MI, Irizarry RA, Kingsford C. Salmon: fast and bias-aware quantification of transcript expression using dual-phase inference. *Nat Methods* 2017;14(4):417–419.

3. Love MI, Huber W, Anders S. Moderated estimation of fold change and dispersion for RNA-seq data with DESeq2. *Genome Biol* 2014;15(12):550.

4. Ritchie ME et al. limma powers differential expression analyses for RNA-sequencing and microarray studies. *Nucleic Acids Res* 2015;43(7):e47–e47.

5. Thompson JJ et al. Blood vessel epicardial substance reduces LRP6 receptor and cytoplasmic β-catenin levels to modulate Wnt signaling and intestinal homeostasis. *Carcinogenesis* 2019;40(9):1086–1098.

6. Thorvaldsdóttir H, Robinson JT, Mesirov JP. Integrative Genomics Viewer (IGV): high-performance genomics data visualization and exploration. *Brief Bioinform* 2013;14(2):178–192.

7. Schaum N et al. Single-cell transcriptomics of 20 mouse organs creates a Tabula Muris. *Nature* 2018;562(7727):367–372.

8. Dieleman et al. Chronic experimental colitis induced by dextran sulphate sodium (DSS) is characterized by Th1 and Th2 cytokines. *Clin Exp Immunol* 1998;114(3):385–391.

9. Fukata M et al. Toll-like receptor-4 promotes the development of colitis-associated colorectal tumors.. *Gastroenterology* 2007;133(6):1869--81.

10. Haber AL et al. A single-cell survey of the small intestinal epithelium. *Nature* 2017;551(7680):333–339.

11. Parikh K et al. Colonic epithelial cell diversity in health and inflammatory bowel disease. *Nature* 2019;567(7746):49–55.

12. Gehart H et al. Identification of Enteroendocrine Regulators by Real-Time Single-Cell Differentiation Mapping. *Cell* 2019;176(5):1158-1173.e16.

13. Li HJ, Ray SK, Kucukural A, Gradwohl G, Leiter AB. Reduced Neurog3 Gene Dosage Shifts Enteroendocrine Progenitor Towards Goblet Cell Lineage in the Mouse Intestine. *Cell Mol Gastroenterology Hepatology* [published online ahead of print: 2020]; doi:10.1016/j.jcmgh.2020.08.006

14. Yan KS et al. Intestinal Enteroendocrine Lineage Cells Possess Homeostatic and Injury-Inducible Stem Cell Activity. *Cell Stem Cell* 2017;21(1):78-90.e6.

15. Böttcher A et al. Non-canonical Wnt/PCP signalling regulates intestinal stem cell lineage priming towards enteroendocrine and Paneth cell fates. *Nat Cell Biol* 2021;23(1):23–31.

16. Kim T-H et al. Broadly permissive intestinal chromatin underlies lateral inhibition and cell plasticity.. *Nature* 2014;506(7489):511 515.

17. Gao S et al. Tracing the temporal-spatial transcriptome landscapes of the human fetal digestive tract using single-cell RNA-sequencing. *Nat Cell Biol* 2018;20(6):721–734.

18. Liberzon A et al. Molecular signatures database (MSigDB) 3.0. *Bioinformatics* 2011;27(12):1739–1740.

19. Powell AE et al. The Pan-ErbB Negative Regulator Lrig1 Is an Intestinal Stem Cell Marker that Functions as a Tumor Suppressor. *Cell* 2012;149(1):146–158.

20. Muñoz J et al. The Lgr5 intestinal stem cell signature: robust expression of proposed quiescent '+4' cell markers. *Embo J* 2012;31(14):3079–3091.

21. Murata K et al. Ascl2-Dependent Cell Dedifferentiation Drives Regeneration of Ablated Intestinal Stem Cells. *Cell Stem Cell* 2020;26(3):377-390.e6.

22. Basak O et al. Induced Quiescence of Lgr5+ Stem Cells in Intestinal Organoids Enables Differentiation of Hormone-Producing Enteroendocrine Cells. *Cell Stem Cell* 2017;20(2):177-190.e4.

23. Basak O et al. Mapping early fate determination in Lgr5+ crypt stem cells using a novel Ki67-RFP allele. *Embo J* 2014;33(18):2057–2068.

24. Smith JR et al. The Year of the Rat: The Rat Genome Database at 20: a multi-species knowledgebase and analysis platform. *Nucleic Acids Res* 2019;48(D1):D731–D742.

25. Kolmykov S et al. GTRD: an integrated view of transcription regulation. *Nucleic Acids Res* 2020;49(D1):gkaa1057-.

26. Wang Y et al. Long-Term Culture Captures Injury-Repair Cycles of Colonic Stem Cells. *Cell* 2019;179(5):1144-1159.e15.

27. Yui S et al. YAP/TAZ-Dependent Reprogramming of Colonic Epithelium Links ECM Remodeling to Tissue Regeneration. *Cell Stem Cell* 2018;22(1):35-49.e7.

28. Mustata RC et al. Identification of Lgr5-Independent Spheroid-Generating Progenitors of the Mouse Fetal Intestinal Epithelium. *Cell Reports* 2013;5(2):421–432.

29. Ayyaz A et al. Single-cell transcriptomes of the regenerating intestine reveal a revival stem cell.. *Nature* 2019;569(7754):121–125.

30. Qu M et al. Establishment of intestinal organoid cultures modeling injury-associated epithelial regeneration. *Cell Res* 2021;1–13.
